# Viscoelasticity in simple indentation-cycle experiments: a computational study

**DOI:** 10.1101/2020.04.15.041640

**Authors:** Yu.M. Efremov, S.L. Kotova, P.S. Timashev

## Abstract

Instrumented indentation has become an indispensable tool for quantitative analysis of the mechanical properties of soft polymers and biological samples at different length scales. These types of samples are known for their prominent viscoelastic behavior, and attempts to calculate such properties from the indentation data are constantly made. The simplest indentation experiment presents a cycle of approach (deepening into the sample) and retraction of the indenter, with the output of the force and indentation depth as functions of time and a force versus indentation dependency (force curve). The linear viscoelastic theory based on the elastic-viscoelastic correspondence principle might predict the shape of force curves based on the experimental conditions and underlying relaxation function of the sample. Here, we conducted a computational analysis based on this theory and studied how the force curves were affected by the indenter geometry, type of indentation (triangular or sinusoidal ramp), and the relaxation functions. The relaxation functions of both traditional and fractional viscoelastic models were considered. The curves obtained from the analytical solutions, numerical algorithm and finite element simulations matched each other well. Common trends for the curve-related parameters (apparent Young’s modulus, normalized hysteresis area, and curve exponent) were revealed. Importantly, the apparent Young’s modulus, obtained by fitting the approach curve to the elastic model, demonstrated a direct relation to the relaxation function for all the tested cases. The study will help researchers to verify which model is more appropriate for the sample description without extensive calculations from the basic curve parameters and their dependency on the indentation rate.

## 1. Introduction

Micro- and nanoindentation (including atomic force microscopy, AFM) have become indispensable tools for the quantitative analysis of the mechanical properties of soft polymers and biological samples with the focus on the corresponding micro- and nanoscale [1–3]. Their benefits include a small sample size and simple preparation, an easily achievable environmental control (e.g. a temperature-controlled fluid cell), a possibility of region-specific mapping and coupling with optical techniques. The purpose of the indentation analysis is to link the indentation data to the meaningful mechanical properties of the sample. Biological samples generally possess time-dependent viscoelastic properties, which can be observed at both the tissue and cellular levels. The important role of viscoelastic properties, as opposed to a purely elastic behavior, has been shown in the studies of different cell phenomena including cancer [4,5], contractile prestress [6], and response to the substrate stiffness [7]. Mathematical models are used to describe the viscoelastic behavior in terms of the relaxation functions. A set of traditional and fractional linear viscoelasticity models are used to describe the sample properties and facilitate the comparison of the parameters across different studies [8–10].

Recently, a large variety of indentation-based methods have been developed to measure and map viscoelastic properties [6,11–14]. Most of them are using modifications of the testing protocol by including constant stress or constant strain phases and oscillatory indentation. However, there are also methods to extract viscoelastic properties directly from the simplest indentation experiment presenting a cycle of approach (deepening into the sample) and retraction of the indenter [5,15–17]. Indentation of a viscoelastic body presents a complex problem with time-varying boundary conditions. The correspondence principle that is used for the problems involving a linear, isotropic viscoelastic body breaks down for the complex indenter shapes and indentation conditions. Lee and Radok obtained an expression for the indentation force and spherical indentation by replacing the elastic modulus in the Hertz solution [18,19] by hereditary integral involving the relaxation response function [20]. However, this solution was found to be invalid when the contact radius decreases with time; the issue was addressed later by Hunter [21], Graham [22] and Ting [23,24]. In any case, the viscoelastic analysis of indentation experiments is much more computationally sophisticated than the analysis based on simple elastic assumptions, and thus, the latter is prevailing in the experimental studies. Moreover, a thorough analysis of force-indentation curves predicted by different viscoelastic models has not been performed before.

This work aims at finding how the viscoelastic relaxation function affects the shape of force curves obtained at different indentation conditions. We analyzed how some basic characteristic curve features change when acquisition parameters such as the indentation rate or indenter shape are varied. These basic features (apparent Young’s modulus, normalized hysteresis area, the curve exponent) can be routinely obtained from experimental curves. Three approaches were used to obtain force curves here: an analytical solution, for the cases where the closed-form analytical solution might be obtained, a numerical solution based on the direct calculation of the Ting’s equations [5], and a FEM simulation-based solution. We performed the analysis for three types of indenter geometries (cylinder, cone, sphere), two types of indentation histories (triangular and sinusoidal ramps), and several types of traditional and fractional viscoelastic models.

## 2. Material and methods

### 2.1. Linear viscoelasticity theory for indentation experiments

We will base the further description on the solution of the indentation problem of a viscoelastic half-space provided by Ting [23,24]. The solution was obtained for the cases of an arbitrary varying radius of the contact area, while here we will concentrate our attention on the load history with a single maximum: the contact area increases first during the approach phase (indenter is pressed into the sample) and then decreases during the retraction phase of the displacement-controlled experiment. The solution for the approach curve coincide with the solution provided by Lee and Radok [20], while the solution for the retraction curve requires an auxiliary function *t*_1_ (*t*). The *t*_1_ (*t*) auxiliary function was introduced as a time point *t*_1_ during the approach phase which corresponds to the same contact area at a time point *t* during the retraction phase. The Lee-Radok’s and Ting’s solutions match for both the approach and retraction curves for a cylindrical indenter and used indentation histories due to the constant contact area. The solution also assumes that a rigid indenter is smooth and axisymmetric but otherwise might have an arbitrary shape. Here we will consider the three most widely used indenter geometries (Fig. 1A): cylinder, sphere, and cone (or pyramid, the difference will be only in the geometrical factor). For the indentation displacement-controlled experiment, the Ting’s solution could be presented in the following form [5]:

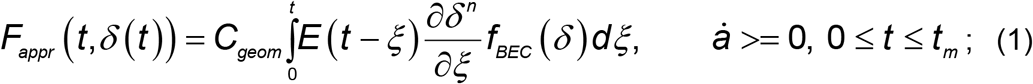

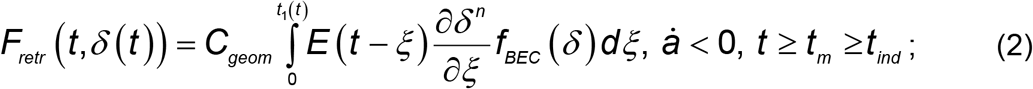

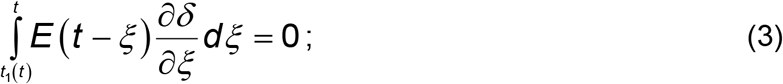

where *F* is the force acting on the cantilever during the approach (*F_appr_*) or the retraction (*F_retr_*); *δ* is the indentation depth; ***a*** is the contact area; *t*_1_(*t*) function is determined by the equation (3); *t* is the indentation time initiated at initial contact (*t_m_* is the time when the maximum contact radius is reached, *t_ind_* is the duration of a complete indentation cycle); *ξ* is the dummy time variable required for the integration; *E*(*t*) is the relaxation function (Young’s relaxation modulus); *n* and *C_geom_* are constants related to the indenter shape [19], e.g.: *n* = 1, *C_cyl_* = 2*R_cyl_*/ (1−*v*^2^) for a cylindrical punch (*R_cyl_* is the radius of the cylinder); *n* = 2, *C_cone_* =2(*tanα_cone_*) / *π* / (1−*v*^2^) for a conical indenter (*α_cone_* is the included half-angle of the cone); *n* =3 / 2, 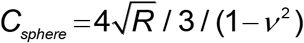 for a paraboloid/spherical indenter (*R* >> *δ* is the radius of the sphere); *v* is the Poisson’s ratio of the sample (assumed to be time-independent). *f_BEC_* (*δ*) is the tip geometry dependent correction coefficient for the finite thickness of the sample attached to the hard substrate [25,26]. This factor does not depend on time and could be neglected when the sample thickness is much larger than the tip-sample contact area. Eq. (1) for *F_appr_* (Lee-Radok’s solution) can also be used to describe the force during the retraction phase, but with limited accuracy. The Ting’s solution requires calculation of *t*_1_(*t*) for the retraction phase. As can be seen from Eq. (3), the *t*_1_(*t*) function is common for all the indenter geometries but depends on the indentation history and the viscoelastic model.

**Figure 1.**
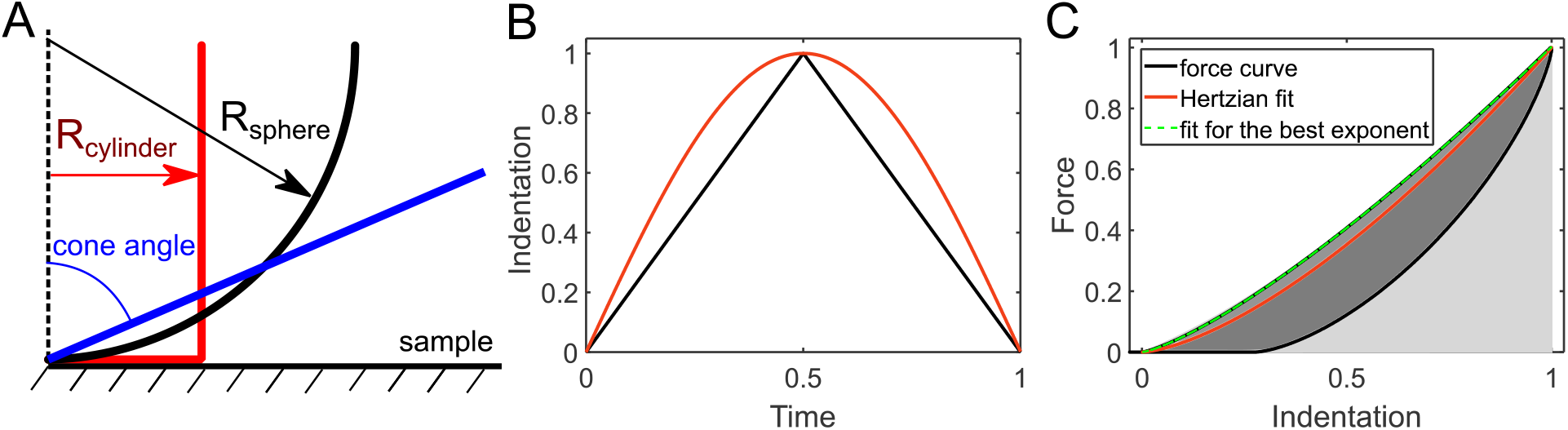
(A) The used indenter (probe) geometries: flat-ended cylinder, sphere, and cone. (B) The applied indentation histories, triangular (black) and sinusoidal (red) ramps. (C) The parameters, extracted for the force curves: apparent Young’s modulus (YM), obtained from the Hertzian fit (red curve); the approach curve exponent, obtained from the fit with an exponent as a fitting parameter; the normalized hysteresis area (NHA), obtained as the area enclosed in the force curve (dark-grey area) divided by the area under the approach curve (light-grey plus dark-grey area). Here, the case of the spherical probe and the springpot viscoelastic model is shown as an example.

The Young’s relaxation modulus *E*(*t*) is a function which determines the viscoelastic behavior of the material and is defined by a specific constitutive viscoelastic model. It is related to the shear stress relaxation modulus as *G*(*t*) = *E*(*t*) / 2 / (1+ *v*) [27]. Note, that the time-independent Poisson’s ratio is assumed here. A “reduced” form of the relaxation (creep) function can be obtained that represents the function normalized by its value at a certain time point, usually, *t* = 0 or *t* = ∞. However, such an approach does not always work since some relaxation functions have infinite or zero values at these time points.

Most modern indenters offer both load and displacement control indentation testing. For example, in a typical AFM experiment, the load is applied by expanding the piezo. The system controls the rate of expansion/retraction of the piezo, but neither force nor indentation histories are controlled directly. The indentation depth *δ* is related to the piezo displacement *Z* as:

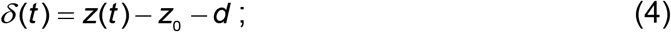

where *z*_0_ is the contact point position (position of the undisturbed sample surface) and *d* is the cantilever deflection. The simplified relation might be considered if the cantilever (or another force sensor) is quite stiff and its deflection is much lower than the indentation depth. If we also place the zero of the displacement axis at the contact point position (*z*_0_=0), the simplified indentation function will be: *δ*(*t*) = *z*(*t*). From here, it is possible to obtain analytical solutions for certain relaxation functions and indentation histories, and this simplification was used in this work.

Here, we will consider two indentation histories, a triangular linear ramp (ramp) and sinusoidal (sin) probe movement (Fig. 1B):

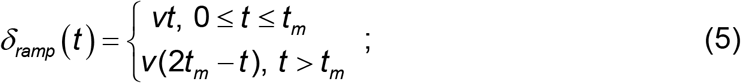

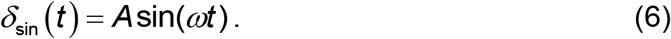

For a triangular linear ramp, the maximum contact radius is reached at *t* = *t_m_* and then it decreases during retraction, the maximum indentation depth is *δ*_*m*_ = *vt_m_*. For the sinusoidal displacement, the maximum contact radius is reached at 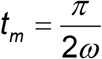 corresponding to a quarter period of the sin wave, the amplitude *A* is set equal to *δ*_*m*_. The *t_m_* value is related to the total probe-sample contact time (indentation time, *t_ind_*) as *t_m_* < *t_ind_* < 2*t_m_*, since the contact area is always present during the approach phase, but would vanish at some point during the retraction phase. The sinusoidal ramp could be beneficial at high indentation rates, since, unlike the triangular ramp, it does not produce an abrupt change in the indentation speed around the turning point. The sinusoidal ramp is used, for example, in the Peak-Force Tapping technique [28] that allows acquisition of force maps with a high speed in AFM experiments.

### 2.2. Numerical and analytical solutions of the Ting’s model

The MATLAB code based on the previous works [5,29] was used here to obtain a numerical solution of the Ting’s model. It calculates the force versus time and the force versus displacement dependencies via numerical differentiation and integration steps, both for the approach and retraction parts of a force curve, the latter involves the numerical calculation of the *t*_1_(*t*) function by an iterative procedure. Arbitrary relaxation functions (in the form of the Young’s relaxation modulus) and indentation histories (currently, with a single maximum in the contact radius versus time data) might be used as an input.

The analytical solutions were obtained for specific viscoelastic models as described in Appendix A. The Python version of the code for the numerical simulation of indentation curves is available at https://github.com/yu-efremov/ViscoIndent.

### 2.3. Finite element analysis

The finite element method (FEM) analysis was performed using the Abaqus CAE software (version 14, Simulia Corp., Providence, RI). The axisymmetric system was created with a cylindrical sample, having a radius of 100 µm and a height of 40 µm. The probe was modeled as a rigid body with the geometries of a flat-ended cylinder (radius of 0.4 µm), sphere (radius of 5 µm), or cone (half-angle of 85°). The sample mesh was optimized for each indenter geometry for a balance between the computational time and accuracy. The probe displacement was assigned for triangular and sinusoidal ramps. The viscoelastic behavior of samples was assigned via the Prony series coefficients. For the power-law rheology model, the relaxation function was approximated as the Prony series expansion including six terms, the coefficients were fitted in MATLAB.

## 3. Results and discussion

The main purpose of the study is to find how the viscoelastic relaxation function of a sample will be reflected in the shape of force curves obtained by indentation. Especially, we are interested in some characteristic features and how they might change when the acquisition parameters such as the indentation rate (inverse of indentation time) are varied. The force curves were obtained using three approaches: an analytical solution, for the cases where the closed-form analytical solution might be obtained, a numerical solution based on the direct calculation of the Ting’s equations, and a FEM simulation solution. The latter one is also a numerical solution by nature, but here we will call it a “simulation solution” to distinguish from the former one. We performed such analysis for three types of indenter geometries (cylinder, cone, sphere), two types of indentation histories (triangular and sinusoidal ramps), and several types of viscoelastic models (Fig. 1).

For all the scenarios (a combination of the probe geometry, indentation history, and viscoelastic model) that provided the closed-form analytical solution, we found a perfect agreement between the analytical and numerical solutions (Fig. S1, S2). This confirms that the numerical solution could effectively substitute the analytical one, and it is especially useful for the cases where the analytical solution could not be obtained. These cases include both complex indentation histories (e.g. non-linear indentation due to a cantilever deflection during the piezo movement, non-linearity in the piezo movement itself) and viscoelastic models (with a large number of elementary elements). The available analytical solutions are presented in Appendix A.

The FEM analysis was performed on two selected sets of parameters for each viscoelastic model, probe geometry and indentation history (total of 24 simulations). The FEM simulation solutions were close to the analytical (where they were obtained) and numerical solutions for all the selected model parameters (Fig. S1, S2). Some observed differences could originate from the finite-size effects. However, the FEM solution is much more time-consuming in comparison with the numerical solution used here.

Therefore, to facilitate the analysis of the force curves, we used only the numerical solutions in the consequent study. Several considerations were taken into account to optimize the analysis:

1. The geometrical parameters of the probe (e.g. cylinder radius, cone angle) will not affect the shape of the curve after the normalization, thus they were not varied.
2. The indentation depth does not affect the shape of the curve since the materials are assumed to behave within the limits of the linear viscoelasticity. Therefore, the change in the indentation depth equals to the change in the indentation speed in the normalized coordinates 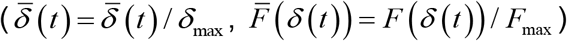.
3. The following parameters were extracted from the force curves. The normalized hysteresis area (NHA), defined as the area enclosed between the approach and retraction curves divided by the area under the approach curve (Fig 1C). This parameter represents the energy dissipation during the indentation cycle, and thus it is especially useful. The second parameter is the power-law exponent of the simple power-law fit applied to the approach curve. Basically, it represents how far the curve deviates from the Hertzian fit. Additionally, the apparent Young’s modulus (YM) extracted from the Hertzian fit of the complete curve (with the fixed exponent value corresponding to the probe geometry) was extracted and its dependency from the indentation time was studied. The graphs are shown in the coordinates of the indentation time *t_ind_*, defined as the total time of indentation cycle, and the corresponding indentation rate is the inverse value of the indentation time.

### 3.1 Simplest spring-dashpot combinations

We begin the analysis with the simplest analytic viscoelastic constitutive models which present a single spring or a dashpot element and their combinations. A spring element symbolizes an ideal elastic behavior; the stress is linearly proportional to the strain: *σ*(*t*) = *kε*(*t*). For this element, the relaxation function is constant in time (*E(t)=E*), and the Ting’s equation solution corresponds to the well-known Hertzian solution of the form:

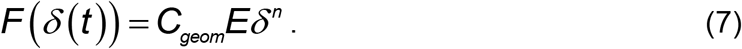

The *n*=1, 1.5, 2 for the cylindrical, spherical and conical probe respectively, the geometrical coefficients are: *C_cylinder_* = 2*R_c_*, 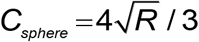, *C_cone_* =2(*tanα_cone_*) / *σ*. The numerical algorithm and FEM simulations provide the force-indentation curves which are analogous to the analytical solution. The force curves show a zero hysteresis area (NHA=0), the curve exponents match with the predicted ones (Fig. 2A).

**Figure 2.**
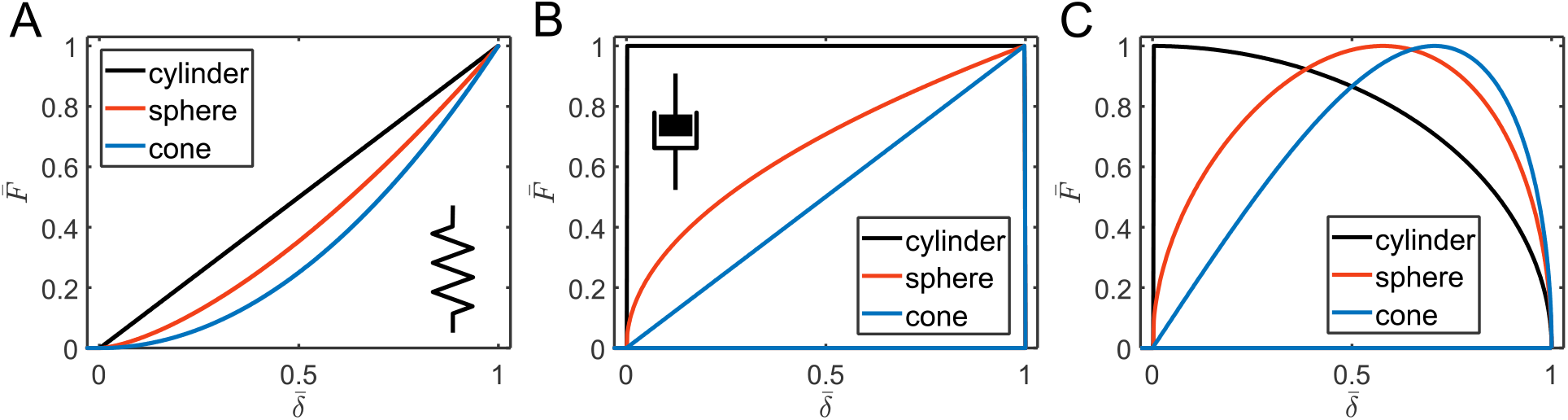
The force versus indentation curves in the normalized coordinates for the spring (A) and dashpot (B, C) elements. The spring element provides the curves described by the Hertzian mechanics, with the curve exponent defined by the probe geometry and zero hysteresis. In contrast, the dashpot provides complete curves with complete hysteresis (zero force during the retraction) that depend on the indentation history, triangular (B) or sinusoidal (C) ramp.

For a dashpot element, the stress is proportional to the strain rate by Trouton’s (or Newton’s) law: *σ*(*t*) =*ηdε*(*t*) / *dt* (*η* is the viscoelastic coefficient or viscosity) according to the behavior of an ideal Newton liquid. The relaxation function is *E*(*t*) = *ηδ_D_*(*t*), where *δ_D_*(*t*) is the Dirac delta function. This viscosity is mostly related to the compressive viscoelastic coefficient (also known as the Trouton coefficient) because indentation measurements involve application of compressive forces normal to the sample surface [30]. The analytical solution (eq. XAppendix) shows that the force drops to zero then the cantilever goes up (retracts), as expected for the viscous material. Thus, the NHA is always equal to one (all energy is dissipated). The shape of the curve is very different from the Hertzian shape and shows a power-law exponent that is lower by one, therefore the Hertzian fit does not provide reasonable data (Fig. 2B, C). Unlike in case of the spring element, the curves now depend on the indentation history and differ for the triangular and sinusoidal ramps. The case of a single dashpot element might correspond to the viscous flow or complete plastic deformation.

The combination of a spring and a dashpot in parallel, known as the Kelvin-Voight element, has the following relaxation function:

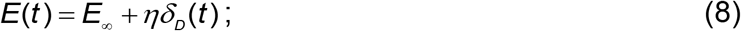

where the subscript “∞” symbolizes a long-term response here and thereafter (E∞ corresponds to the long-term modulus). The characteristic time of the model is 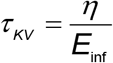. At short indentation times *t_ind_* << *τ*_*KV*_, the behavior is dominated by the dashpot, and at long indentation times *t_ind_* >> *τ_KV_* – by the spring (the spring modulus corresponds to *E*_∞_). Accordingly, the NHA decreases with the indentation time, it is close to zero at slow rates, and close to one at fast rates (Fig. 3). The effective YM is proportional to the indentation rate at short times, but then it reaches a plateau corresponding to *E*_∞_ at long times. The curve exponent at long times is equal to the Hertzian one, but as for the single dashpot model, it is lower by one at short times. In the actual experiments, such a huge deviation from the Hertzian exponent could indeed be observed when the dissipation (NHA) is large, for example in AFM experiments on cells at very high indentation rates [29] (Fig. S3). A notable feature of the model is the jump in the force at the initial contact for the case of the cylindrical indenter, that is caused by the dashpot and could be seen in FEM simulations as well [25] (Fig. S1). The Kelvin-Voight element can describe the plastic flow of the material during the indentation. In the normalized coordinates 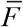 versus 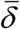, the curve shape is determined solely by the *t_ind_* / *τ_KV_* ratio. The curves of NHA, YM, and exponent versus normalized contact time only weakly depend on the probe geometry (Fig 3C). The data for the sinusoidal ramp demonstrated similar trends and are not shown here.

**Figure 3.**
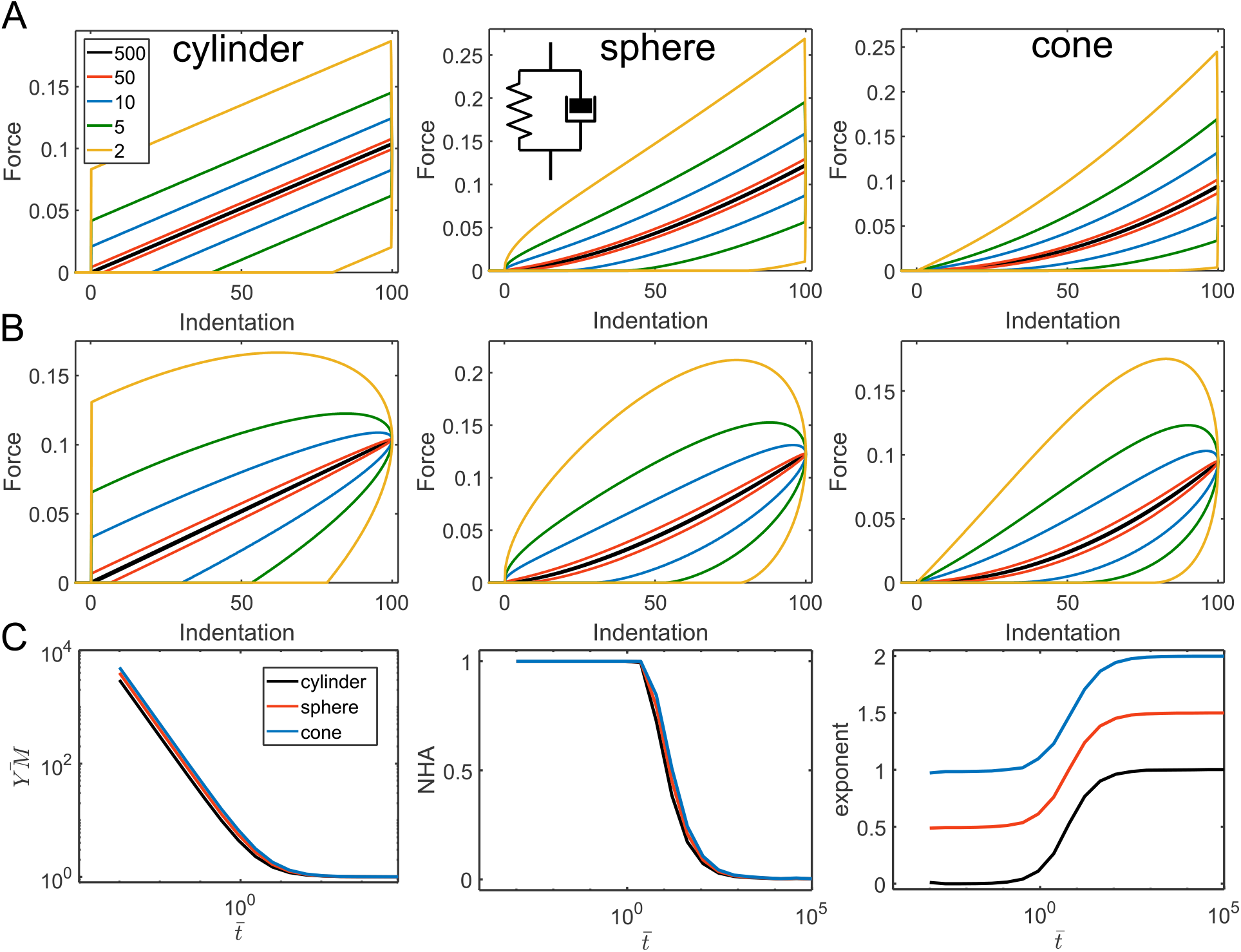
The force curves and parameters acquired from the force curves for the Kevin-Voight model. (A) The force curves for the triangular displacement; cylindrical, spherical, and conical indenters, and varied *t_ind_* / *τ*_*KV*_ ratio (shown with different line colors). (B) The force curves for the sinusoidal displacement. (C) Dependencies for the normalized YM (*YM* / *E*_∞_), NHA, and curve exponent on the normalized indentation time 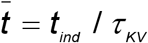.

For another combination, a spring and a dashpot in series, known as the Maxwell element, the relaxation function presents a well-known exponential decay:

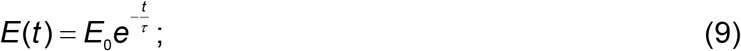

where the subscript “0” symbolizes the instantaneous response here and thereafter (spring modulus corresponds to the instantaneous modulus *E*_0_). The characteristic time 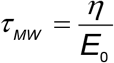. At short indentation times *t_ind_* << *τ*_*MW*_ the behavior is dominated by the spring, and at long indentation times *t_ind_* >> *τ_MW_* – by the dashpot. Accordingly, the NHA increases with the indentation time, it is close to zero at fast rates, and close to one at slow rates, which is opposite to the Kelvin-Voight model. The effective YM is proportional to the indentation rate at long times, but a plateau corresponding to *E*_0_ is observed at short indentation times. The curve exponent at short times is equal to the Hertzian one and is lower by one at long times. Again, the probe geometry and ramp type only weakly affect the observed dependencies (Fig. 4).

**Figure 4.**
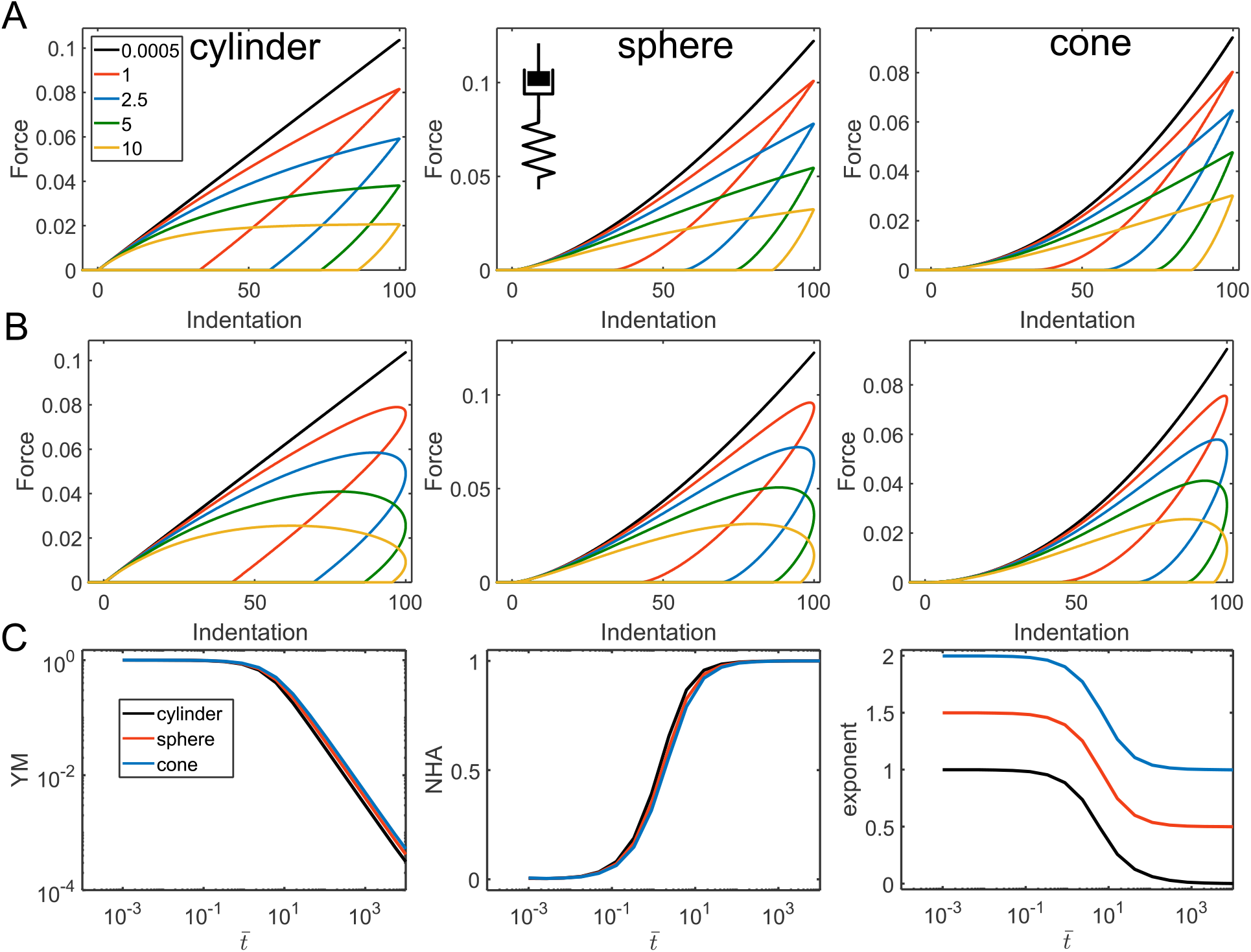
The force curves and parameters acquired from the force curves for the Maxwell model. (A) The force curves for the triangular displacement; cylindrical, spherical, and conical indenters, and varied *t_ind_* / *τ_MW_* ratio (shown with different line colors). (B) The force curves for the sinusoidal displacement. (C) Dependencies for the normalized YM (*YM* / *E*_0_), NHA, and curve exponent on the normalized indentation time 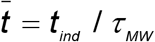.

The standard linear solid model (Zener model) can be represented as the Maxwell element in parallel with a second spring (*E*_*s*2_) that determines the long-term modulus of the system. The short-term modulus is a combination of the moduli of the two springs *E*_0_ = *E*_*s*1_ + *E*_*s*2_. The relaxation function is:

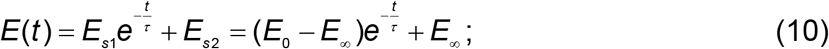

which differs from the relaxation function of the Maxwell element by the presence of the *E*_∞_ term. The model has the characteristic times 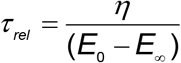 and 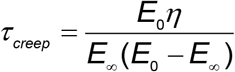, known to be characteristic times of relaxation and creep, respectively. From here, for the larger *E*_0_ / *E*_∞_ ratio and the same *τ_rel_* value, the relaxation will be more pronounced and will take more time. At short and long indentation times, there are plateaus for all dependencies, with the moduli corresponding to *E*_0_ and *E*_∞_, respectively, the curve exponent corresponding to the Hertzian one, and NHA close to zero. The maximum NHA is observed at values slightly larger than *τ_rel_* and *τ_creep_*; the larger *E*_0_ / *E*_∞_ ratio provides a larger and wider negative peak. At the very large *E*_0_ / *E*_∞_ ratios, the model behaves as a single dashpot in this intermittent regime, while at small ratios the viscoelastic behavior will be unnoticeable. The curve exponent is affected in a similar way (Fig. 5A-C).

**Figure 5.**
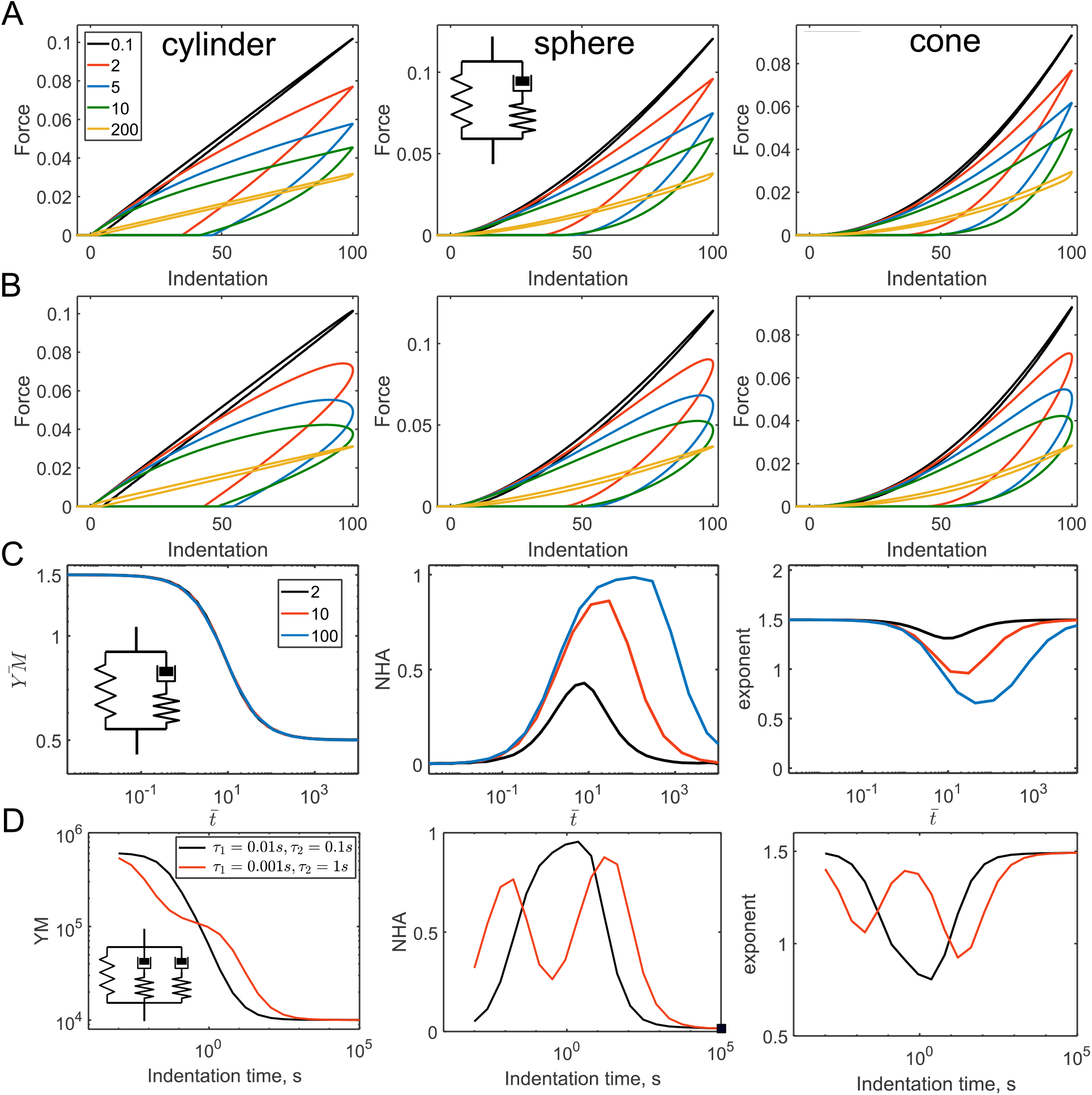
The force curves and parameters acquired from the force curves for the SLS model. (A) The force curves for the triangular displacement; cylindrical, spherical, and conical indenters, and varied *t_ind_* / *τ_rel_* ratio (shown with different line colors). (B) The force curves for the sinusoidal displacement. (C) Dependencies for the normalized YM (*YM* / (*E*_0_ − *E*_∞_)), NHA, and curve exponent on the normalized indentation time 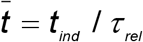 for different *E*_0_ / *E*_∞_ ratios. (D) Generalized Maxwell model with two relaxation times, two cases with a small and large difference in the relaxation times. Dependencies for the YM, NHA, and curve exponent on the indentation time.

The SLS model could be seen as a particular case of the generalized Maxwell model, where several Maxwell elements are connected in parallel. We analyzed a case with two such elements and a spring:

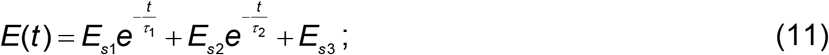

where *E*_0_ = *E*_*s*1_ + *E*_*s*2_ + *E*_*s*3_, *E*_inf_ = *E*_*s*3_. The main outcome of such a model is that, locally, near the relaxation time of one of the Maxwell elements, the shape of the force curves will be determined by this particular element. If the relaxation times of both Maxwell elements are close to each other, then larger and wider peak for the NHA and curve exponent will be observed instead of two separate peaks (Fig. 5D).

### 3.2. Fractional viscoelastic models

From the Fractional Calculus theory, another type of the basic viscoelastic element is a so-called springpot with a governing equation *σ*(*t*) = *K*_*α*_*d*^*α*^*ε* (*t*) / *dt^α^*. The element has two parameters, the unitless power-law exponent and the parameter *K_α_* with units of [Pa s^−α^]. The Young’s relaxation function can be written as:

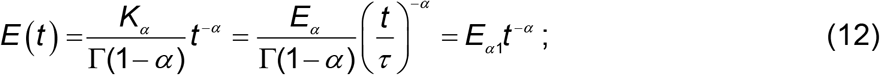

where Γ() is the Gamma function. It is worth stressing the meaning of the other parameters in the equation. The *K_α_* parameter with units of [Pa s^-α^] is not very convenient and does not have a straightforward physical meaning. It is commonly replaced with *K*_*α*_ = *E*_*α*_ *τ*^*α*^, where *E_α_* is the Young modulus in [Pa] and *τ* is in [s]. However, these two parameters are not independent and are linked via *K_α_*. To reduce the number of independent parameters back to two, we might assign *τ*==1 s, then the parameter *E*_*α*1_ = *E*_*α*,*τ*=1_ / Γ(1− *α*) will arise with units of [Pa] and a simple meaning of the value of the relaxation function at t=1s. However, for the correspondence of the units, the time in the last part of Eq. (12) should be considered as unitless (*t* / [*τ*= 1*S*]). Notably, the model can be easily rescaled to any other characteristic time *τ*.

The springpot element intermediates between a spring and a dashpot through a fractional-order derivative *α* of the strain history (0<*α* <1). The force curves constructed with the model demonstrate some interesting features: 1) In the normalized coordinates, the shape of the curve is defined solely by *α* ; 2) The NHA and curve exponent parameters are independent of the indentation time and are also determined by *α*. The larger *α* value corresponds to the larger hysteresis. The curve exponent value approximately equals to its Hertzian value minus *α* ; 3) Effective Young’s modulus increases with the indentation rate following the power-law dependency with the same exponent *α* (Fig. 6C). These effects were similar for all the considered geometries, for the sinusoidal and triangular ramp loadings. Such behavior makes a simple guidance for the identification of the power-law behavior in experiments: the constant hysteresis in the force curves acquired at different rates, the power-law dependency of the effective modulus on the indentation rate. Such effects are indeed observed in experiments on cells in a wide range of indentation rates [5] (Fig. S4).

**Figure 6.**
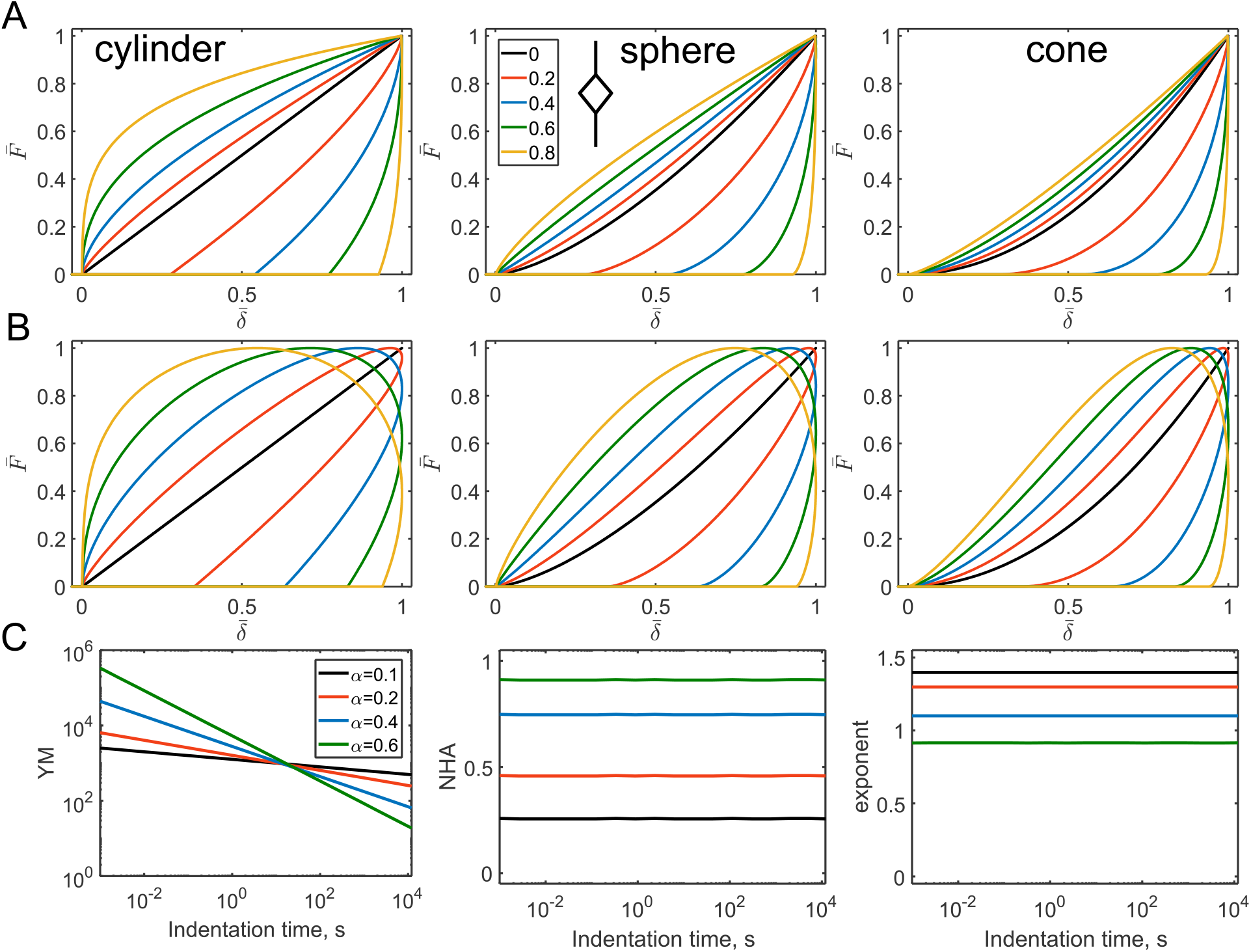
The force curves and parameters acquired from the force curves for the springpot model. (A) The force curves for the triangular displacement; cylindrical, spherical, and conical indenters, and varied *α*. (B) The force curves for the sinusoidal displacement. (C) Dependencies for the YM, NHA, and curve exponent on the indentation time, only for the spherical indenter, triangular displacement.

The springpot element could be used in various combinations with other elements, including other springpot elements. We will address some basic combinations here. A springpot in parallel with a spring will provide the following relaxation function:

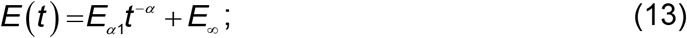

where *E*_∞_ < *E*_*α*1_. This combination is also known as the fractional Kelvin-Voight model [31,32]. The difference with a single dashpot element is the presence of the long-term modulus at slow indentation rates. Therefore, the YM, curve exponent and NHA observe the gradual prolonged transition between the power-law and elastic regimes. The transition point is defined by the characteristic time 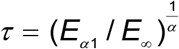 [s]; at shorter times, the behavior is dominated by the springpot. Thus, hysteresis increases with the indentation rate toward the certain limit defined by the *α* value (Fig. 7).

**Figure 7.**
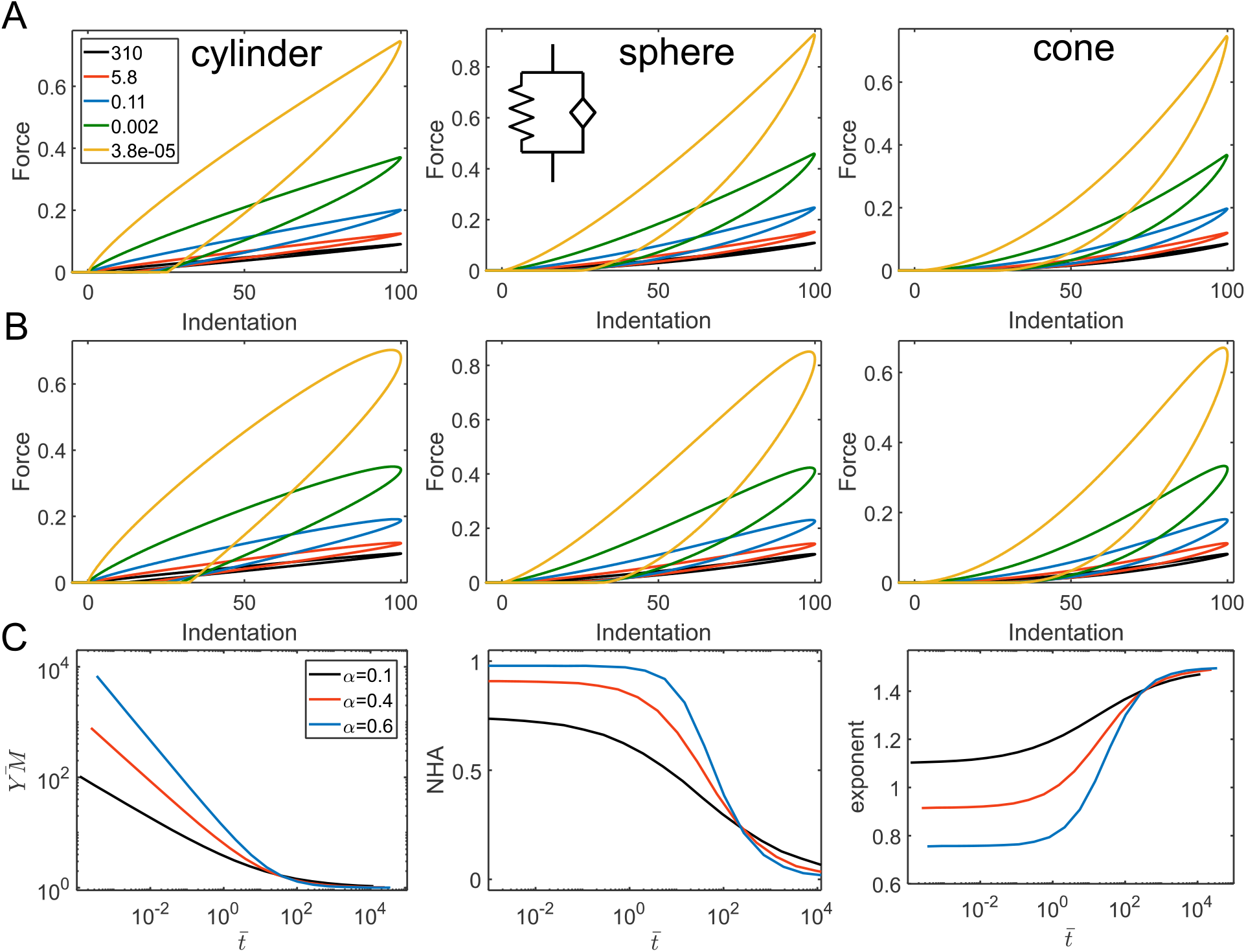
The force curves and parameters acquired from the force curves for the fractional Kelvin-Voight model. (A) The force curves for the triangular displacement; cylindrical, spherical, and conical indenters, *α* = 0.2 and varied *t_ind_*/ *τ* ratio. (B) The force curves for the sinusoidal displacement. (C) Dependencies for the normalized YM, NHA, and curve exponent on the normalized indentation time 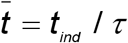 for different *α* values.

For a springpot and a spring in series, the resulted relaxation function is more complex due to the presence of the Mittag-Lefler function (ML) in the equation:

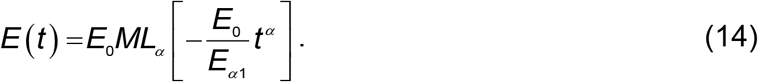

The Mittag-Leffler function is a special function that arises from the solution of certain fractional differential equations and is calculated numerically. As opposed to the parallel combination, now the spring (*E_s_* = *E*_0_) dominate in response at short timescales and the springpot – at long timescales (Fig. 8). The characteristic transition time is *E*_0_ / *E*_*α*1_ [s].

**Figure 8.**
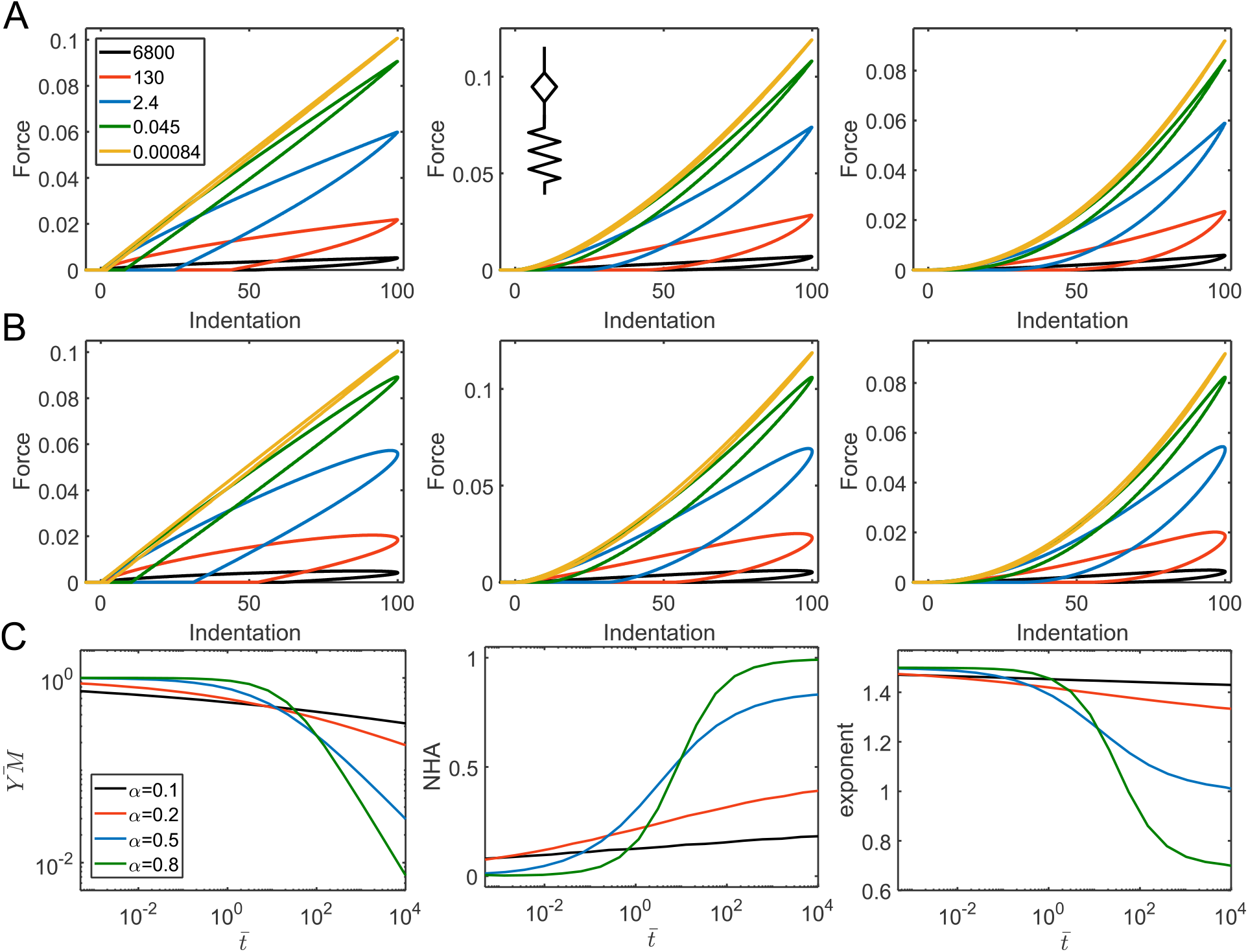
The force curves and parameters acquired from the force curves for the model representing by a springpot and a spring in series. (A) The force curves for the triangular displacement; cylindrical, spherical, and conical indenters, *α* = 0.4 and varied *t_ind_* / *τ* ratio. (B) The force curves for the sinusoidal displacement. (C) Dependencies for the normalized YM, NHA, and curve exponent on the normalized indentation time 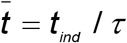 for different *α* values.

The same analogy could be observed in combinations of a springpot and a dashpot: when placed in parallel, the short-time response is controlled by the dashpot, and when in series – by the springpot. The relaxation functions are:

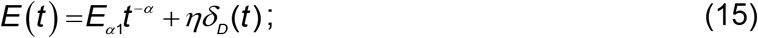

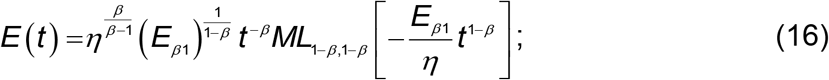

for the parallel and serial arrangements, respectively (Fig. 9, A-B). More generally, when a combination of two springpot elements is considered, the parallel arrangement leads to the short-time behavior controlled by the element with the higher exponent, and the long-time response – by the element with the lower exponent. The opposite is true when the elements are arranged in series. The relaxation functions for two springpot elements are as follows:

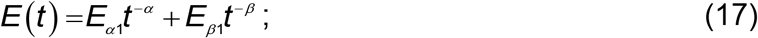

for the parallel arrangement; and for the serial arrangement:

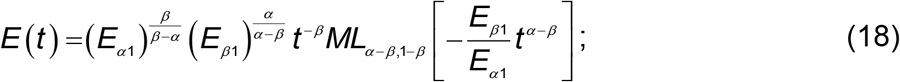

where *α* > *β* (Fig. 9, C-D). All the previously considered models for serial arrangements of the elements could be seen as a particular case of the two serial springpot elements. When *α* = 1, the springpot element reduces to the dashpot, and when *β* = 0, it reduces to the spring. The characteristic transition times, as was shown in [8], can be presented as 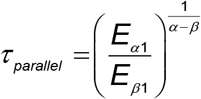 and 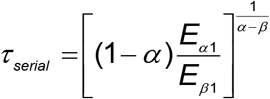.

**Figure 9.**
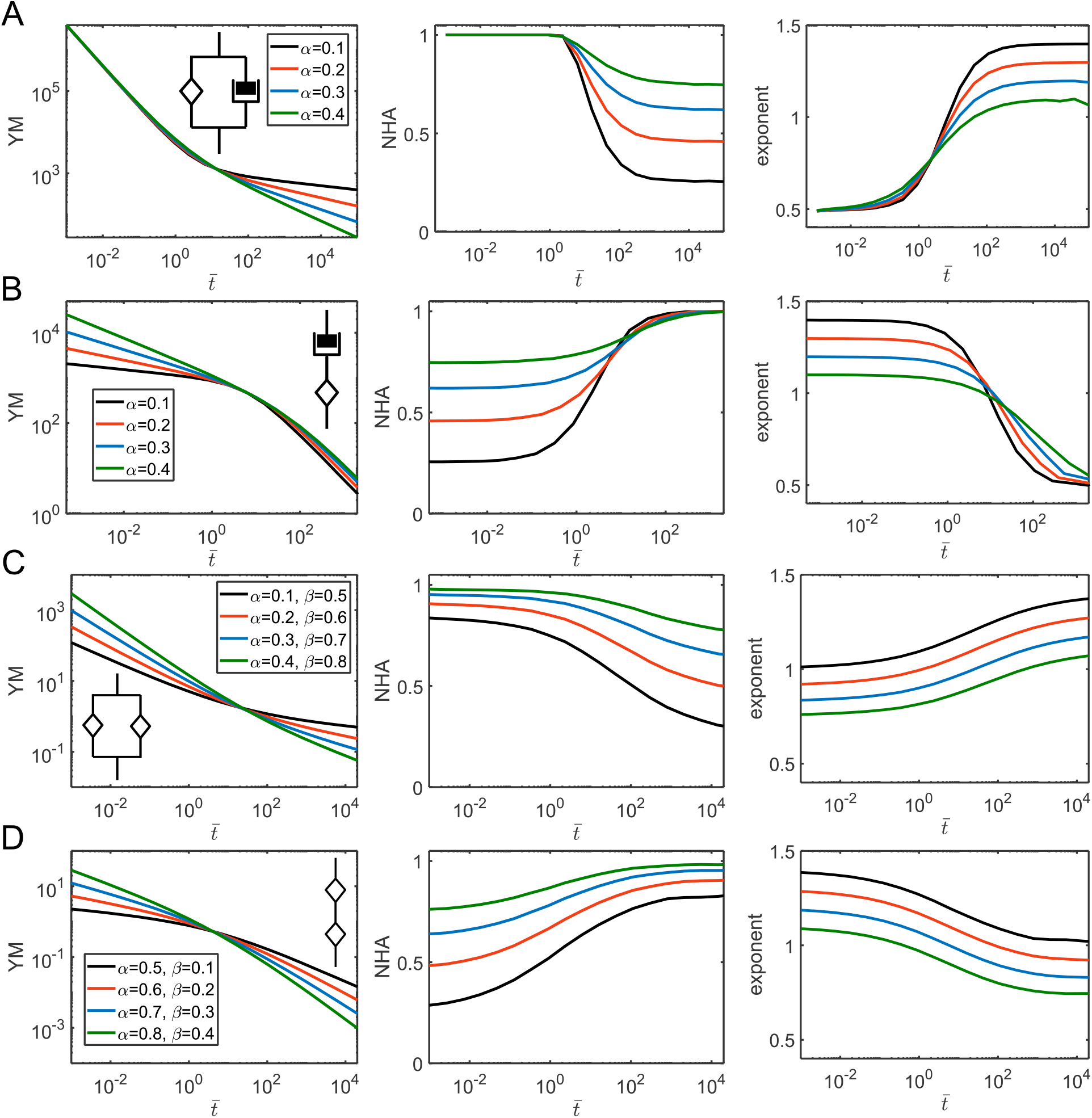
The parameters acquired from the force curves for a springpot and a dashpot in different combinations. Dependencies for the YM, NHA, and curve exponent on the normalized indentation time 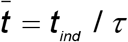, where *τ* is the characteristic time of the corresponding model, for different *α* (and *β*) values for a springpot in parallel with a dashpot (A); a springpot in series with a dashpot (B); a springpot in parallel with another springpot; (D) a springpot in series with another springpot.

The addition of a spring in parallel to another element results in the addition of the long-term modulus *E*_∞_ to the relaxation function. We will consider two such models here, the first one is a spring in parallel with a springpot and spring combination, also known as a fractional SLS (fractional Zener) model [33], with the following relaxation function:

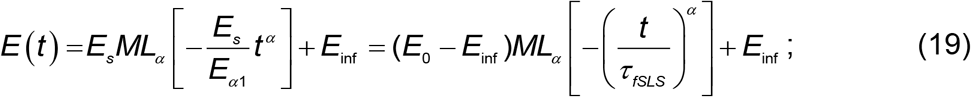

where *E*_0_ = *E_s_* + *E*_inf_ and the characteristic time is 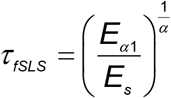. As in the common SLS model, this model has plateau regions at short and long times corresponding to *E*_0_ and *E*_inf_ elastic moduli. At *α* = 0, the model reduces to the SLS model. Increase in *α* leads to the stretching of the transition region around the transition time, similar to the model known as the stretched exponent model [34]. Accordingly, the hysteresis in force curves is observed over a wider range of indentation times, while diminishing to zero at extremes (Fig. 10A).

**Figure 10.**
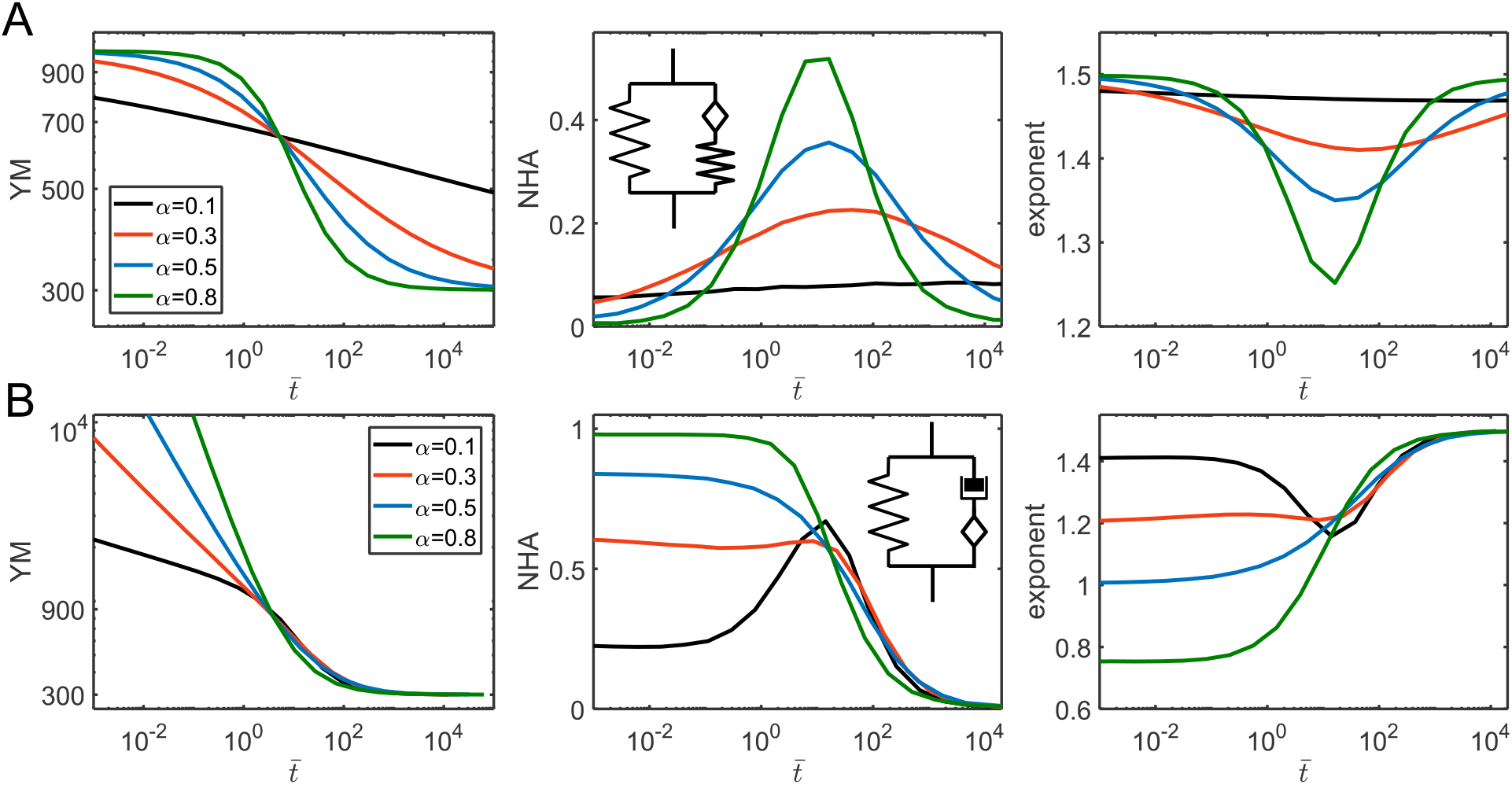
The parameters acquired from the force curves for the two three-element fractional viscoelastic models: a spring in parallel with a springpot and spring combination (A), a spring in parallel with a springpot and dashpot combination (B). Dependencies for the YM, NHA, and curve exponent on normalized indentation time 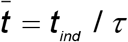, where *τ* is the characteristic time of the corresponding model, for different *α* values.

The second model is a spring in parallel with a springpot and dashpot combination. The model was comprehensively studied in the recent works [8,35]. The relaxation function is:

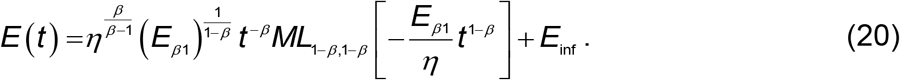

The behavior of the model, as expected, is similar to the springpot - dashpot combination with a transition to the elastic regime at the long timescales. On the other hand, the model reduces to the SLS model when *β* = 0. The largest hysteresis in force curves is observed in the middle region where the dashpot is active, then, at short timescales, the hysteresis reduces toward the values defined by the springpot exponent *β* (Fig. 10B).

### 3.3 General dependencies of the curve parameters on the relaxation function

Some general dependencies could be drawn from the provided numerical analysis. Importantly, the shape of the force curves is strongly affected by the relaxation function of the material. A huge deviation from the Hertzian curve exponent (up to minus one) is possible then relaxation is significant. It correlates with the large hysteresis, NHA, of the curve. Since the elastic assumption is often used to fit the force curves, the lower curve caused by viscoelasticity might be misinterpreted as non-linear elasticity (strain-softening), since the fit of shallower regions will provide a higher YM and the fit of the deeper regions – a lower YM. Otherwise, if the real non-linearity like strain-stiffening is presented, viscoelasticity might conceal it to some extent.

The dependencies of the apparent YM, NHA, and curve exponent versus indentation time were close for the studied indenter geometries and indentation histories. By comparison with the actual relaxation functions, we have found that the apparent YM corresponds very closely to the time-averaged value of the relaxation function with the limits from *t* = 0 to *t* = *t_ind_* /4 (Fig. 11A):

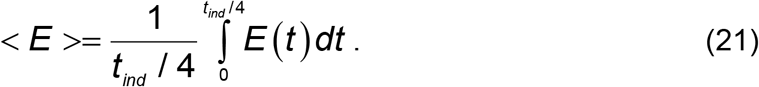

**Figure 11.**
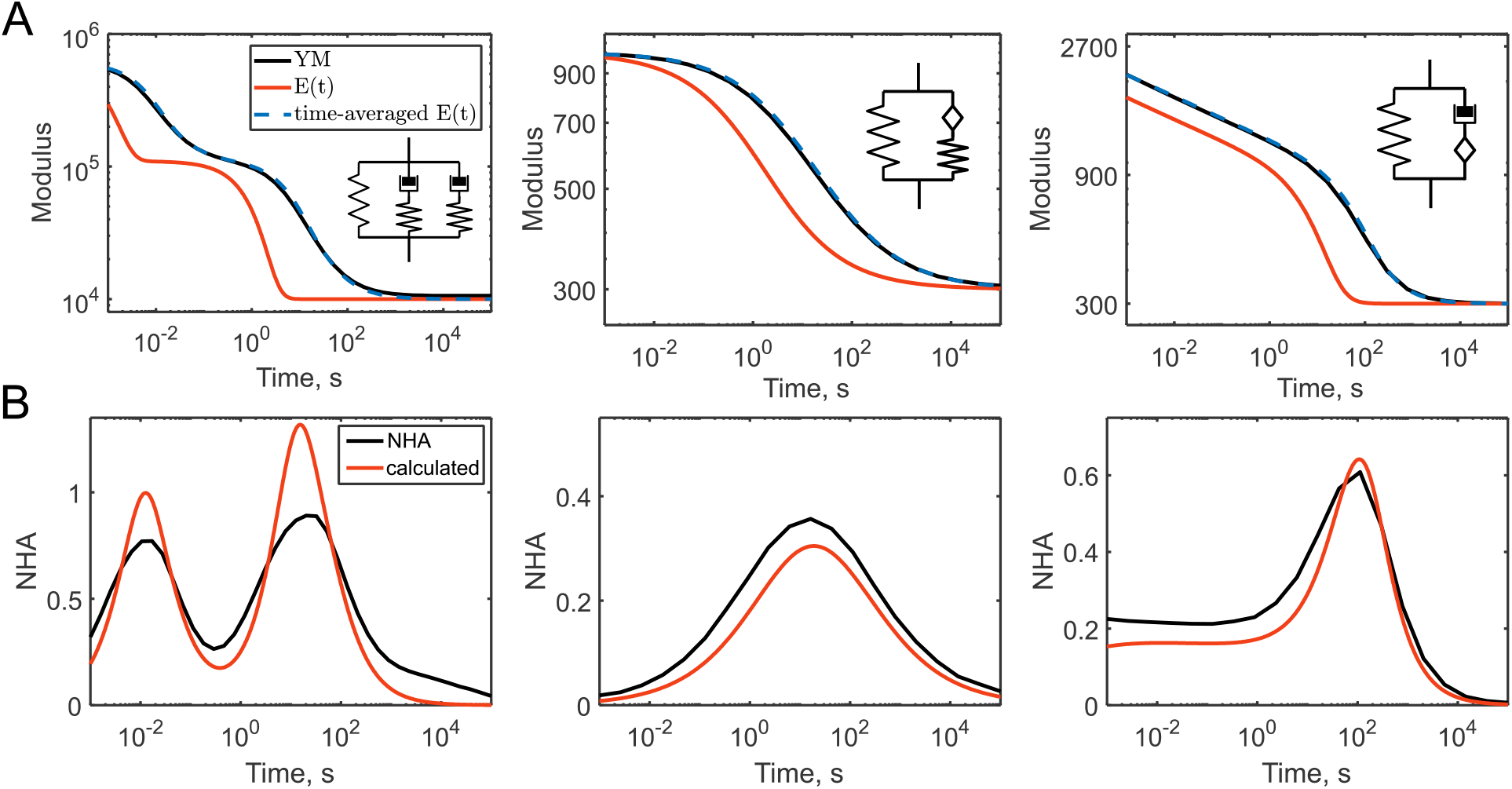
Comparison of the apparent YM (A) and NHA (B) acquired from the force curves with the relaxation function for the three viscoelastic models: Generalized Maxwell model with two relaxation times; a spring in parallel with a springpot and spring combination; a spring in parallel with a springpot and dashpot combination. (A) the YM, the relaxation function *E(t)*, and the time-averaged relaxation function. (B) the NHA and slope of the time-averaged relaxation function on the logarithmic scale.

The upper limit corresponds to half of the approach time since only the approach curve data are used for the YM calculation. The dependencies of the NHA and curve exponent values are more complex. As a first approximation, they are related to the local slope of the relaxation function on a logarithmic scale. Indeed, we have found that the doubled slope of the time-averaged relaxation function, shown in Fig. 11A, is close to the observed NHA values (Fig. 11B). A zero slope corresponds to the elastic regions with a zero NHA and Hertzian exponent value; and the maximum slope, which is equal to one (for the dashpot), leads to the NHA=1 and a decrease in the exponent by one. It should also be noted, that NHA is a fraction of the energy dissipated during the indentation cycle, and thus it is related to the loss tangent (ratio of the loss modulus to the storage modulus) at the frequency corresponding to 1/ *t_ind_*.

## 4. Concluding remarks

The shape of the force-indentation curves is well-predicted by the underlying viscoelastic relaxation function; the analytical, numerical, and simulation solutions provide well-matched results. Viscoelasticity causes substantial deviation of the curve shape from the purely elastic response. The curve exponent decreases from the Hertzian value, which might be misinterpreted as strain-softening if the elastic assumption is used. The presence of the hysteresis area is a clear sign of the viscoelastic response (in the absence of strong adhesion and plasticity).

Several approaches might be used to extract the information about the relaxation function from the indentation data. One of simple ways is to observe the dependencies of the apparent YM and NHA on the indentation rate and to compare them with the predictions from the relaxation functions. For example, the presence of the constant NHA over the wide range of times is an indicator of the power-law rheology. The idea of using the YM versus indentation rate was implemented in the previous studies [36,37]. In the case of the power-law rheology, the power-law exponent might be extracted from such a dependency. The empirical comparison shows that the apparent modulus does not follow the relaxation function exactly, but it is very close to the time-averaged value of this function (Fig 11A). The NHA is related to the local slope of the relaxation function on a logarithmic scale. A large hysteresis in the curve correlates with the strong dependency of the YM on the indentation rate.

More advanced approaches involve algorithms for the fitting of the force curves to the preselected viscoelastic models [5,15,16] or algorithms for the direct reconstruction of the relaxation function. In both cases, however, it is useful to obtain experimental curves in a wide range of indentation rates, and thus, it is useful to know how the curves should look for different viscoelastic models.

## Acknowledgments

This study was supported by the Russian academic excellence project “5-100” (FEM simulations), by the Russian Science Foundation under the grant No. 19-79-00354 (Yu.M.E., development of numerical algorithms) and by the grant of the President of the Russian Federation for young scientists MK‐1613.2020.7 (Yu.M.E., data analysis).

## Appendix A. Analytical solutions of the Ting’s equations for selected viscoelastic models, probe geometries and indentation histories.

For the acquisition of analytical solutions, the equations were solved symbolically with SageMath [38] and Wolfram Alpha integral calculator [39]. We will reproduce the relaxation functions here and provide analytical solutions for the Ting’s equations describing a common indentation experiment with the triangular or sinusoidal displacement. The solutions for the approach (tip deepens into the sample, contact area increasing) and retraction (contact area decreasing) curves will be provided separately. The solutions for the approach curve (Lee-Radok’s solution) could also be extended for the case of the retraction curves, we did it for some situations where the complete Ting’s solution was not obtained. The Lee-Radok’s and Ting’s solution match for the cylindrical probe since the contact area is constant or has a zero value.

The **spring element**, *σ*(*t*) = *kε*(*t*). The relaxation function is constant in time (*E(t)=E*), and the Ting’s equation solution corresponds to the well-known Hertzian solution of the form:

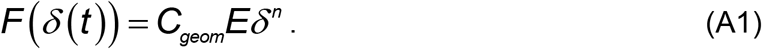

The n=1, 1.5, 2 for the cylindrical, spherical and conical probe, respectively, the geometrical coefficients are: *C_cylinder_* = 2*R_c_* / (1−*v*^2^), 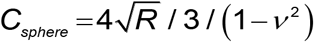, *C_cone_* = 2(*tanα*) / *π* / (1−*v*^2^).

The **dashpot element**, *σ* (*t*) *ηdε*(*t*) / *dt*(*η* is the viscoelastic coefficient or viscosity) according to the behavior of an ideal Newton liquid. The relaxation function is *E*(*t*) = *ηδ_D_*(*t*), where *δ_D_* (*t*) is the Dirac delta function. The common analytical solution of the Ting’s equation for all the geometries is:

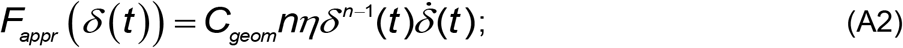

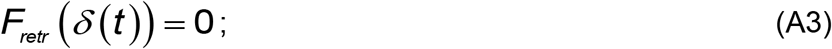

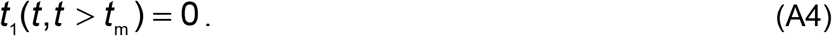

As expected for a viscous material, the force drops to zero then the cantilever goes up (retracts).

The **Kelvin-Voight element**, a combination of a spring and a dashpot in parallel, has the following relaxation function: *E*(*t*) = *E*_∞_ + *ηδ_D_*(*t*). We will split the solutions for the approach *F_ap_* and retraction *F_retr_* curves. *t*_1_(*t*) function can be found from Eq. (3), which leads to the condition:

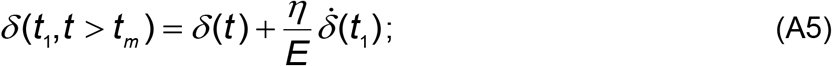

which for the triangular ramp leads to:

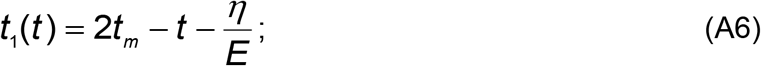

and for the sinusoidal ramp:

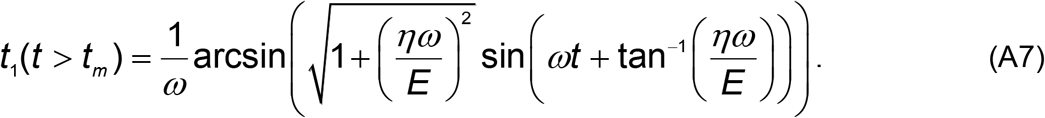

The common solution for all the geometries for the triangular (ramp) and sinusoidal (sin) indentation histories are, respectively:

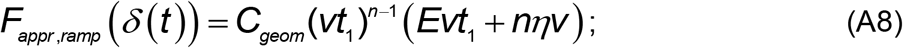

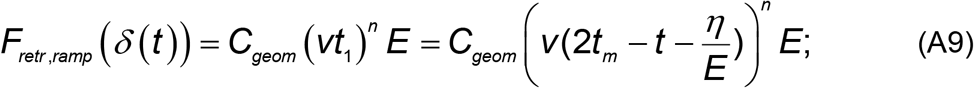

[eq

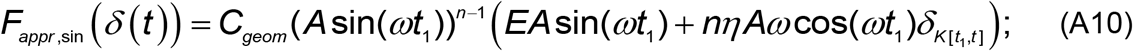

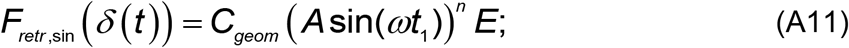

solution for the approach curve was also obtained before in [30].

For the **Maxwell element**, the relaxation function is: 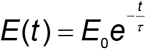. The *t*_1_ (*t*) function for the triangular displacement is:

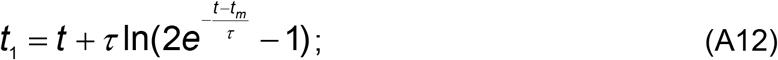

The solution for *t*_1_ (*t*) in the case of the sinusoidal load could not be isolated, but it can be numerically found from the following relation:

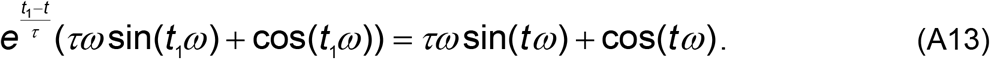

For the approach curve, the solutions for the triangular displacement and different probe geometries are:

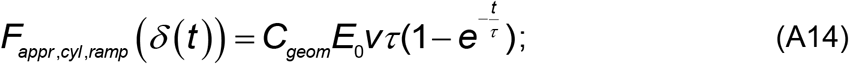

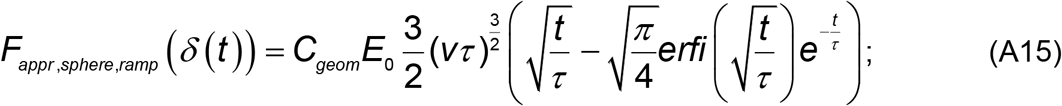

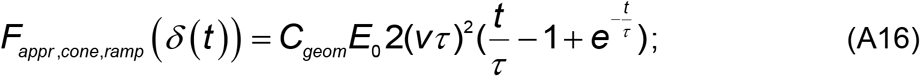

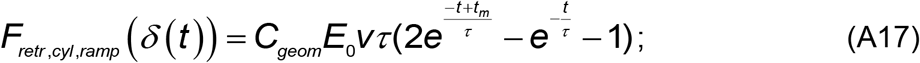

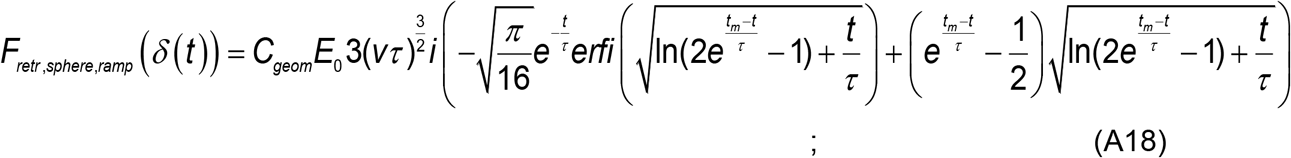

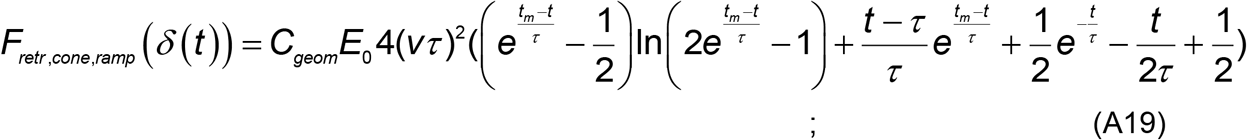

where *erfi()* is the imaginary Gauss error function. For the sinusoidal displacement, there is no closed-form analytical solution for the case of the spherical probe:

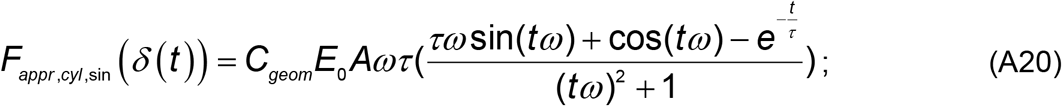

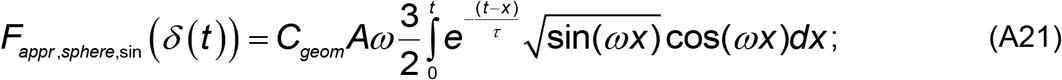

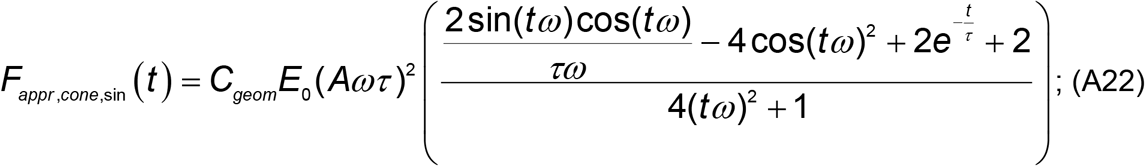

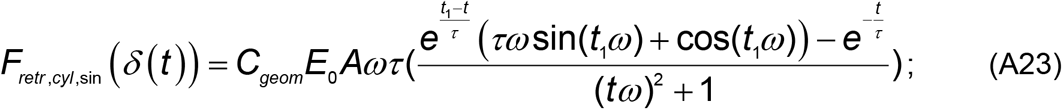

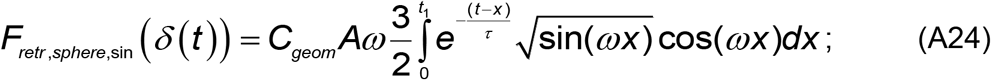

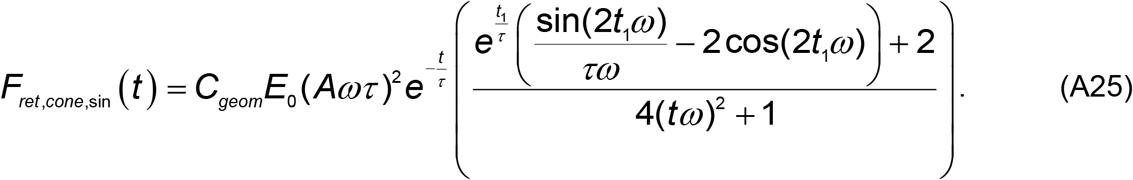

By comparing Eqs. A12-14 and A23, it follows that for the cylindrical indenter the solutions for the approach and retraction curves match, as stated above. It is true for other considered viscoelastic models as well.

The relaxation function of the **Standard linear solid** model is 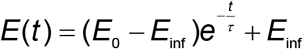. The *t*_1_ (*t*) function for the triangular ramp is:

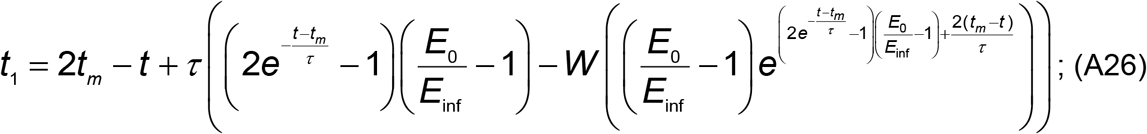

where W is the Lambert W function. For the sinusoidal displacement, the *t*_1_(*t*) can be numerically found from the following relation:

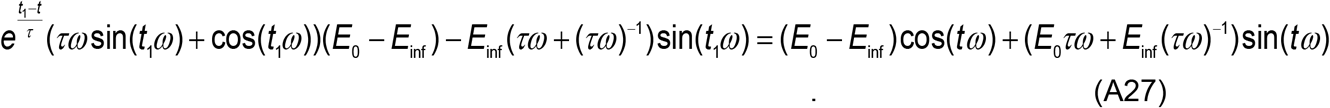

The solutions for the approach curves are:

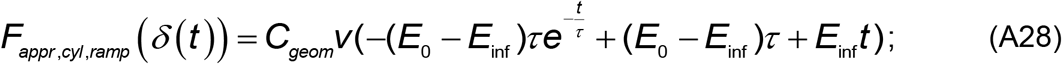

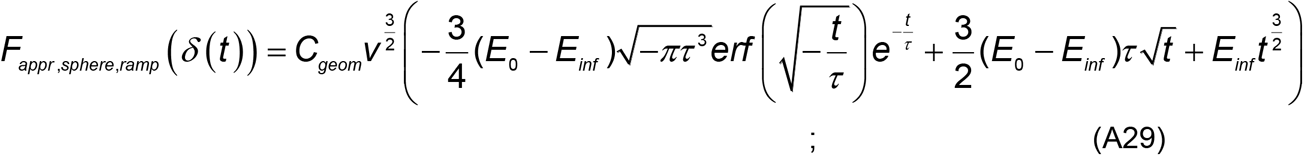

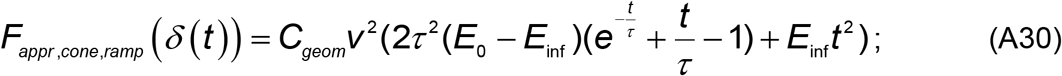

For the retraction curves, the analytical solutions for the spherical and conical probes are too long and complex, so here only the solution for the cylindrical probe is presented:

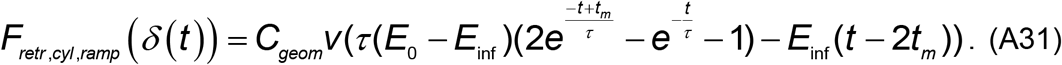

The solutions for the sinusoidal displacement were found except for the case of sphere-retraction, but are not presented here due to complexity.

For a single springpot element (Power-law rheology model), we will use the Young’s relaxation function in the form of *E* (*t*) = *E*_*α*1_ *t*^−*α*^. The analytical solutions for the Ting’s equations for triangular indentation were obtained previously by Bruckner et al. [15]. The *t*_1_(*t*) function for the triangular ramp is:

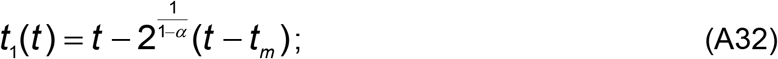

the solution was not acquired for the sinusoidal ramp.

The solutions for the triangular ramp are as follows:

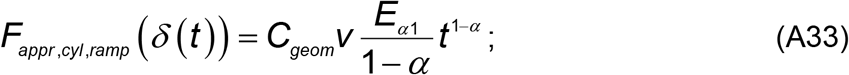

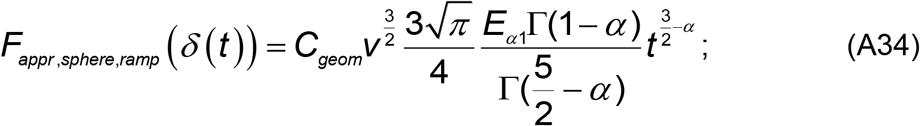

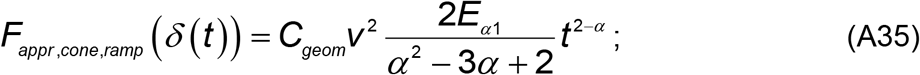

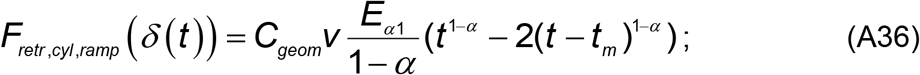

The solutions for the retraction curves for the spherical and conical geometries are too long and not presented here. For the sinusoidal ramp, only the following solutions were obtained for the approach curves:

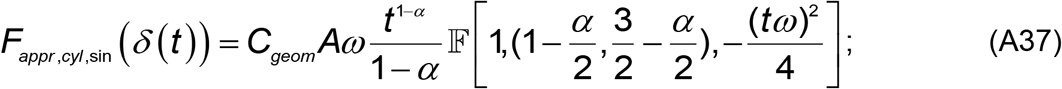

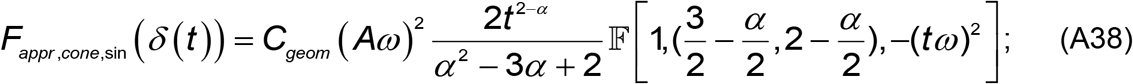

where 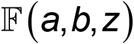 is the generalized hypergeometric function. The analytical solutions could be obtained for some other viscoelastic functions and particular sets of indentation histories and probe geometries, but it is beyond the tasks of the current study.

## Supplementary material

**Figure S1.**
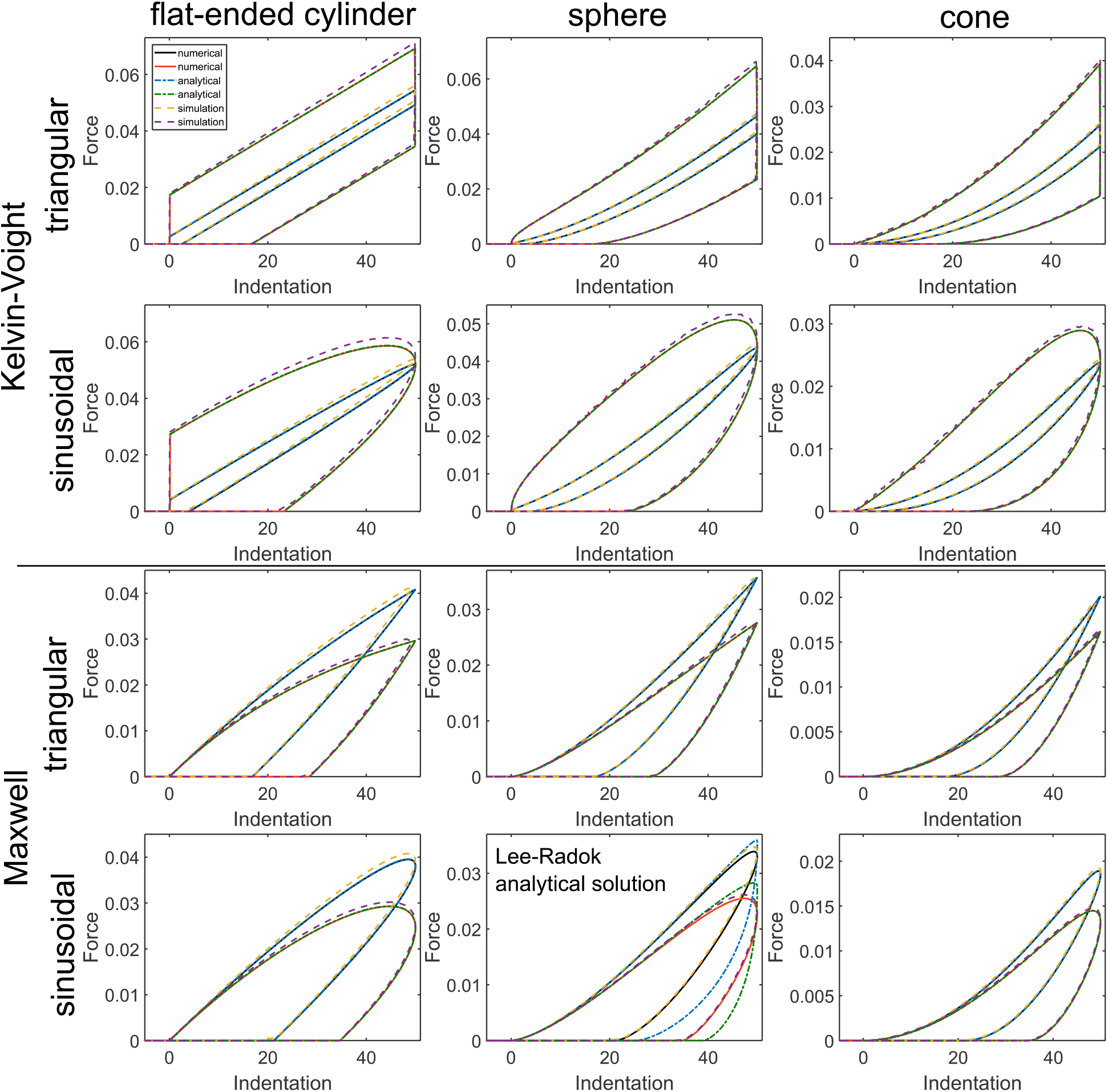
A comparison of numerical, analytical, and simulation solutions for the Kelvin-Voight and Maxwell models, different probe geometries (flat-ended cylinder, sphere, and cone) and indentation histories (triangular and sinusoidal probe displacement). The parameters of the Kelvin-Voight model: 1) *E*_∞_ =1000 Pa, *η* =10 Pa*s; 2) *E*_∞_ =1000 Pa, *η* =100 Pa*s. The parameters of the Maxwell model: 1) *E*_0_ =1000 Pa, *τ*=2 s; 2) *E*_0_ =1000 Pa, *τ* =0.8 s. The indentation speed for the triangular ramp was 50 nm/s, the frequency of the sinusoidal ramp was 0.25 Hz (total time was 2 s for both cases), the amplitude was 50 nm. There is no analytical Ting’s solution for the Maxwell model-sphere-sinusoidal ramp case, the Lee-Radok’s solution is presented.

**Figure S2.**
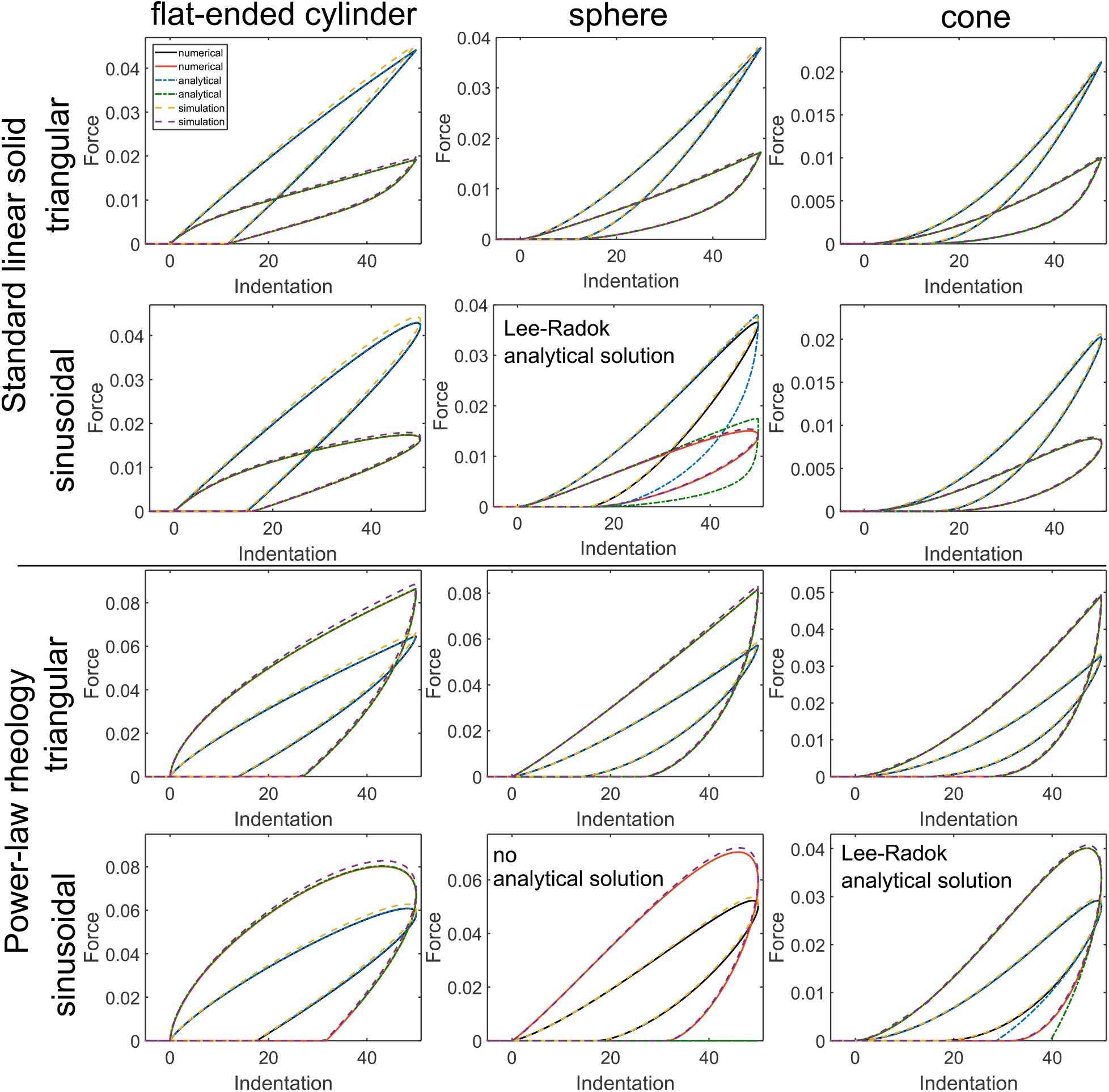
A comparison of numerical, analytical, and simulation solutions for the SLS and PLR models, different probe geometries (flat-ended cylinder, sphere, and cone) and indentation histories (triangular and sinusoidal probe displacement). The parameters of the SLS model: 1) *E*_0_ =1000 Pa, *τ* =2 s, *E*_∞_ =300 Pa; 2) *E*0 =1000 Pa, *τ* =0.5 s, *E*_∞_ =300 Pa. The parameters of the PLR model: 1) *E*_*α*1_ =1000 Pa, =0.2; 2) *E*_0_ =1000 Pa, *α* =0.4. The indentation speed for the triangular ramp was 50 nm/s, the frequency of the sinusoidal ramp was 0.25 Hz (total time was 2 s for both cases), the amplitude was 50 nm. There is no analytical Ting’s solution for the SLS model-sphere-triangular ramp, PLR model-sphere-cone-sinusoidal ramp cases, the Lee-Radok’s solution is presented where available.

**Figure S3.**
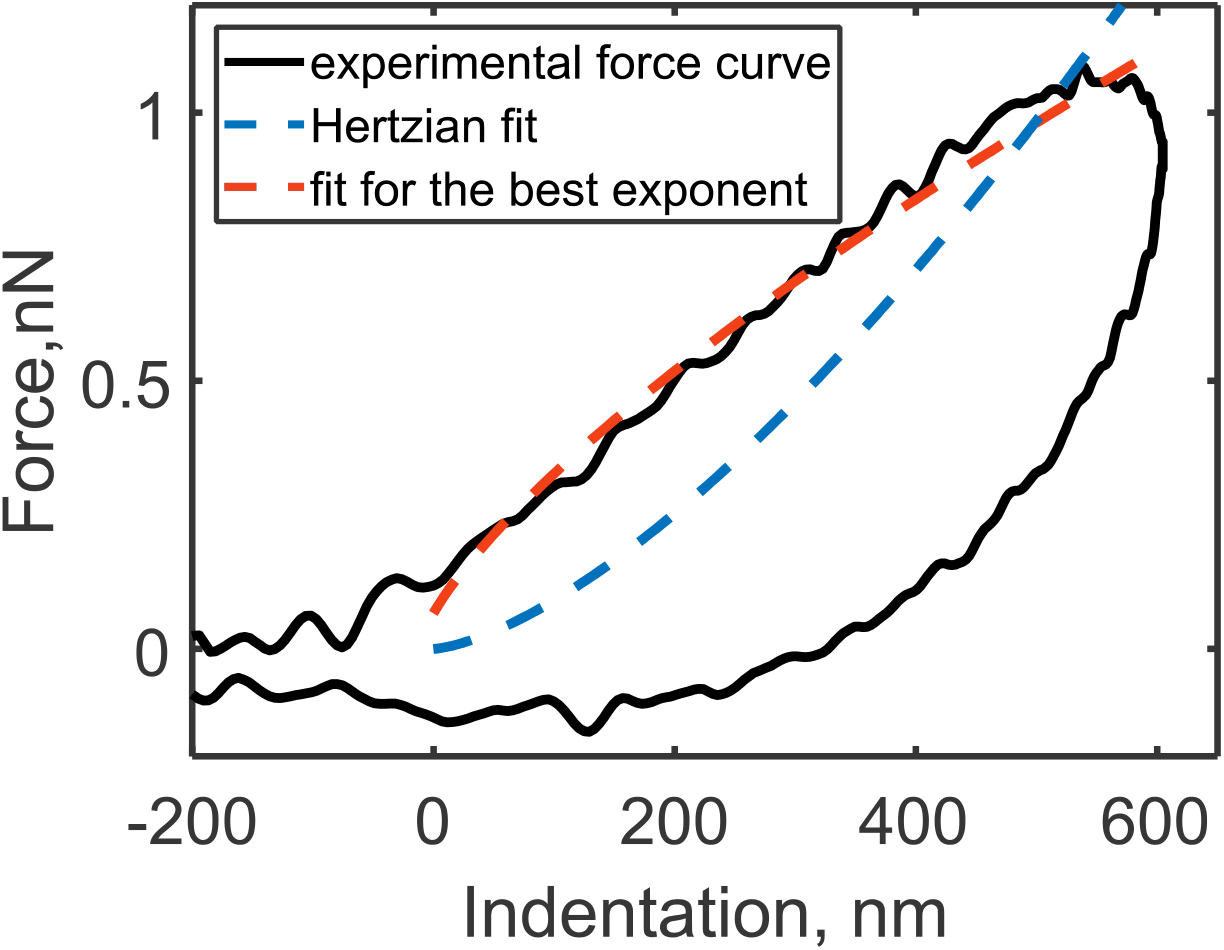
A force curve obtained with AFM indentation on a cell (NIH 3T3 fibroblast) at a high indentation rate. The force curve was obtained at the indentation rate of 660 Hz that corresponds to the indentation time of 0.0015 s, 140 nm diameter spherical (parabolical) probe, sinusoidal displacement. Due to a strong dissipation (the NHA=0.81), the Hertzian fit does not follow the curve closely, and the curve exponent (0.73) is twice lower than the Hertzian one (1.5). The experimental data are taken from [29].

**Figure S4.**
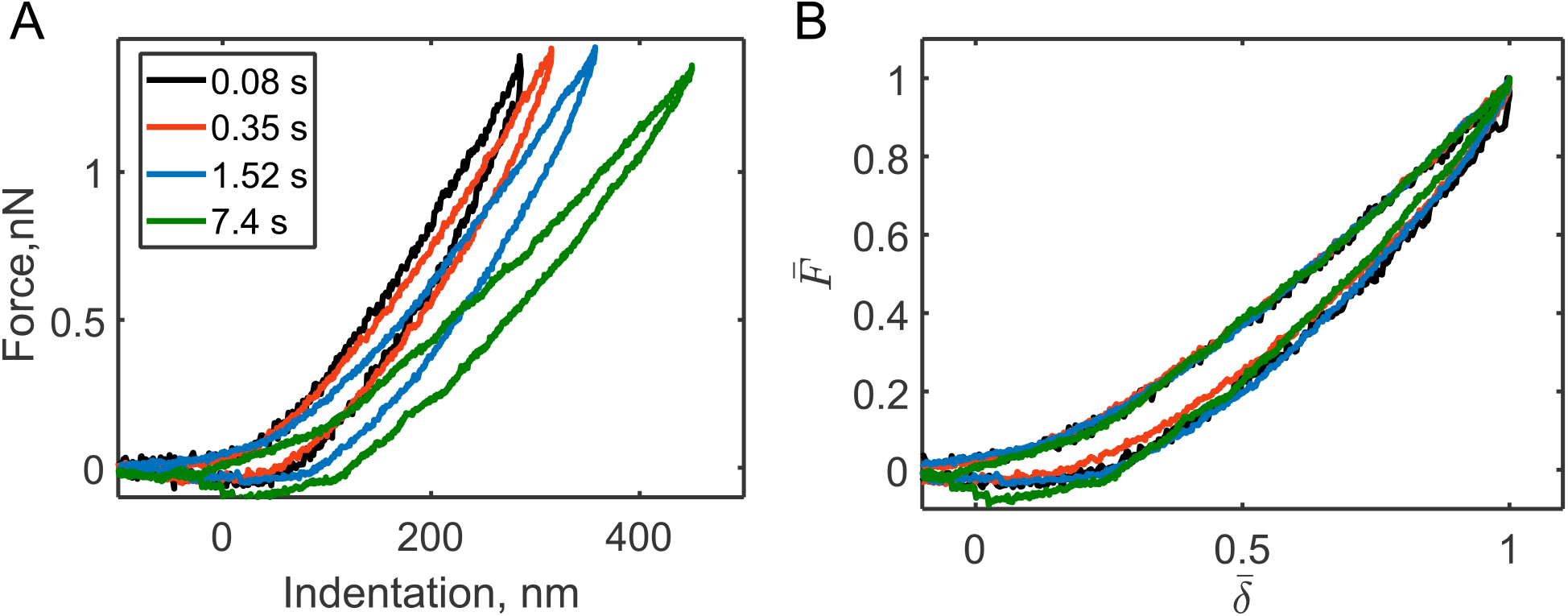
Force curves obtained with AFM indentation on cells (NIH 3T3 fibroblasts) can be described well with the power-law rheology model (single springpot). **(A)** The force curves obtained at different indentation times (showed in the legend) over three orders of magnitude, a 5 μm diameter spherical probe. (B) In the normalized coordinates, the curves match each other well since they all have very close values of the NHA and curve exponent. Moreover, these values are close to the values from the numerical prediction (NHA = 0.28 vs 0.26; curve exponent = 1.4 vs 1.4; for the experimental *α* ≈ 0.1). The experimental data are taken from [5].

## References

[1] L. Qian, H. Zhao, Nanoindentation of soft biological materials, Micromachines. 9 (2018). doi:10.3390/mi9120654.

[2] D.M. Ebenstein, L. a. Pruitt, Nanoindentation of biological materials, Nano Today. 1 (2006) 26–33. doi:10.1016/S1748-0132(06)70077-9.

[3] Y.M. Efremov, T. Okajima, A. Raman, Measuring viscoelasticity of soft biological samples using atomic force microscopy, Soft Matter. 16 (2020) 64–81. doi:10.1039/C9SM01020C.

[4] J. Rother, H. Nöding, I. Mey, A. Janshoff, Atomic force microscopy-based microrheology reveals significant differences in the viscoelastic response between malign and benign cell lines., Open Biol. 4 (2014) 140046. doi:10.1098/rsob.140046.

[5] Y.M. Efremov, W.-H. Wang, S.D. Hardy, R.L. Geahlen, A. Raman, Measuring nanoscale viscoelastic parameters of cells directly from AFM force-displacement curves, Sci. Rep. 7 (2017) 1541. doi:10.1038/s41598-017-01784-3.

[6] N. Schierbaum, J. Rheinlaender, T.E. Schäffer, Combined atomic force microscopy (AFM) and traction force microscopy (TFM) reveals a correlation between viscoelastic material properties and contractile prestress of living cells, Soft Matter. 15 (2019) 1721–1729. doi:10.1039/C8SM01585F.

[7] C. Rianna, M. Radmacher, Comparison of viscoelastic properties of cancer and normal thyroid cells on different stiffness substrates, Eur. Biophys. J. 46 (2017) 309–324. doi:10.1007/s00249-016-1168-4.

[8] A. Bonfanti, J.L. Kaplan, G. Charras, A.J. Kabla, Fractional viscoelastic models for power-law materials, ArXiv. (2020) 1–28. http://arxiv.org/abs/2003.07834.

[9] R. Lakes, Viscoelastic materials, Cambridge University Press, Cambridge, 2009. doi:10.1017/CBO9780511626722.

[10] R.L. Magin, Fractional Calculus in Bioengineering, Part 2, Crit. Rev. Biomed. Eng. 32 (2004) 105–194. doi:10.1615/CritRevBiomedEng.v32.i2.10.

[11] Y.M. Efremov, A.X. Cartagena-Rivera, A.I.M. Athamneh, D.M. Suter, A. Raman, Mapping heterogeneity of cellular mechanics by multi-harmonic atomic force microscopy, Nat. Protoc. 13 (2018) 2200–2216. doi:10.1038/s41596-018-0031-8.

[12] M. Dokukin, I. Sokolov, High-resolution high-speed dynamic mechanical spectroscopy of cells and other soft materials with the help of atomic force microscopy, Sci. Rep. 5 (2015) 12630. doi:10.1038/srep12630.

[13] R. Takahashi, T. Okajima, Mapping power-law rheology of living cells using multi-frequency force modulation atomic force microscopy, Appl. Phys. Lett. 107 (2015) 173702. doi:10.1063/1.4934874.

[14] Y.T. Cheng, F. Yang, Obtaining shear relaxation modulus and creep compliance of linear viscoelastic materials from instrumented indentation using axisymmetric indenters of power-law profiles, J. Mater. Res. 24 (2009) 3013–3017. doi:10.1557/jmr.2009.0365.

[15] B.R. Brückner, H. Nöding, A. Janshoff, Viscoelastic Properties of Confluent MDCK II Cells Obtained from Force Cycle Experiments, Biophys. J. 112 (2017) 724–735. doi:10.1016/j.bpj.2016.12.032.

[16] J.S. de Sousa, J.A.C. Santos, E.B. Barros, L.M.R. Alencar, W.T. Cruz, M. V. Ramos, J. Mendes Filho, Analytical model of atomic-force-microscopy force curves in viscoelastic materials exhibiting power law relaxation, J. Appl. Phys. 121 (2017) 034901. doi:10.1063/1.4974043.

[17] M. V. Ramesh Kumar, R. Narasimhan, Analysis of spherical indentation of linear viscoelastic materials, Curr. Sci. 87 (2004) 1088–1095.

[18] H. Hertz, Über die Berührung Fester Elastischer Körper, J. Für Die Reine u. Angew. Math. 92 (1881) 156–171. http://www.citeulike.org/group/13900/article/11394774 (accessed April 25, 2013).

[19] I.N. Sneddon, The relation between load and penetration in the axisymmetric Boussinesq problem for a punch of arbitrary profile, Int. J. Eng. Sci. 3 (1965) 47–57. http://www.sciencedirect.com/science/article/pii/0020722565900194 (accessed August 14, 2014).

[20] E.H. Lee, J.R.M. Radok, The Contact Problem for Viscoelastic Bodies, J. Appl. Mech. 27 (1960) 438–444.

[21] S.C. Hunter, The Hertz problem for a rigid spherical indenter and a viscoelastic half-space, J. Mech. Phys. Solids. 8 (1960) 219–234. doi:10.1016/0022-5096(60)90028-4.

[22] G.A.C. Graham, The contact problem in the linear theory of viscoelasticity when the time dependent contact area has any number of maxima and minima, Int. J. Eng. Sci. 5 (1967) 495–514. doi:10.1016/0020-7225(67)90037-7.

[23] T.C.T. Ting, The contact stresses between a rigid indenter and a viscoelastic half-space, J. Appl. Mech. 33 (1966) 845–854. doi:10.1115/1.3625192.

[24] T.C.T. Ting, Contact Problems in the Linear Theory of Viscoelasticity, J. Appl. Mech. 35 (1968) 248. doi:10.1115/1.3601188.

[25] P.D. Garcia, R. Garcia, Determination of the viscoelastic properties of a single cell cultured on a rigid support by force microscopy, Nanoscale. 10 (2018) 19799–19809. doi:10.1039/C8NR05899G.

[26] P.D. Garcia, R. Garcia, Determination of the Elastic Moduli of a Single Cell Cultured on a Rigid Support by Force Microscopy, Biophys. J. 114 (2018) 2923–2932. doi:10.1016/j.bpj.2018.05.012.

[27] L.D. Landau, E.M. Lifshitz, Theory of Elasticity, Pergamon Press, 1989.

[28] S.B. Kaemmar, Introduction to Bruker’s ScanAsyst and PeakForce Tapping AFM Technology, Appl. Note. 133 (2011) 12. https://www.bruker.com/fileadmin/user_upload/8-PDF-Docs/SurfaceAnalysis/AFM/ApplicationNotes/Introduction_to_Brukers_ScanAsyst_and_PeakForce_Tapping_Atomic_Force_Microscopy_Technology_AFM_AN133.pdf.

[29] Y.M. Efremov, A.I. Shpichka, S.L. Kotova, P.S. Timashev, Viscoelastic mapping of cells based on fast force volume and PeakForce Tapping, Soft Matter. 15 (2019) 5455–5463. doi:10.1039/C9SM00711C.

[30] P.D. Garcia, C.R. Guerrero, R. Garcia, Time-resolved nanomechanics of a single cell under the depolymerization of the cytoskeleton, Nanoscale. 9 (2017) 12051–12059. doi:10.1039/C7NR03419A.

[31] H. Zhang, Y. Wang, M.F. Insana, Ramp-hold relaxation solutions for the KVFD model applied to soft viscoelastic media, Meas. Sci. Technol. 27 (2016) 25702. doi:10.1088/0957-0233/27/2/025702.

[32] H. Zhang, Q.Z. Zhang, L. Ruan, J. Duan, M. Wan, M.F. Insana, Modeling ramp-hold indentation measurements based on Kelvin–Voigt fractional derivative model, Meas. Sci. Technol. 29 (2018) 035701. doi:10.1088/1361-6501/aa9daf.

[33] B. Carmichael, H. Babahosseini, S.N. Mahmoodi, M. Agah, The fractional viscoelastic response of human breast tissue cells, Phys. Biol. 12 (2015) 46001. doi:10.1088/1478-3975/12/4/046001.

[34] T. Okajima, M. Tanaka, S. Tsukiyama, T. Kadowaki, S. Yamamoto, M. Shimomura, H. Tokumoto, Stress relaxation of HepG2 cells measured by atomic force microscopy, Nanotechnology. 18 (2007) 084010. doi:10.1088/0957-4484/18/8/084010.

[35] A. Bonfanti, J. Fouchard, N. Khalilgharibi, G. Charras, A. Kabla, A unified rheological model for cells and cellularised materials, R. Soc. Open Sci. 7 (2020) 190920. doi:10.1098/rsos.190920.

[36] M.A. Caporizzo, C.M. Roco, M.C.C. Ferrer, M.E. Grady, E. Parrish, D.M. Eckmann, R.J. Composto, Strain-rate Dependence of Elastic Modulus Reveals Silver Nanoparticle Induced Cytotoxicity, Nanobiomedicine. 2 (2015) 9. doi:10.5772/61328.

[37] Y.M. Efremov, A.A. Dokrunova, D. V Bagrov, K.S. Kudryashova, O.S. Sokolova, K. V Shaitan, The effects of confluency on cell mechanical properties, J. Biomech. 46 (2013) 1081–1087. doi:10.1016/j.jbiomech.2013.01.022.

[38] The Sage Developers, SageMath, the Sage Mathematics Software System (Version 8.7), (2019). https://www.sagemath.org.

[39] W. Research, WolframAlpha integral calculator, (2019). https://www.wolframalpha.com/calculators/integral-calculator.

